# Interpreting Convolutional Neural Networks in Population Genetics

**DOI:** 10.64898/2025.12.04.692404

**Authors:** Huiting Xu, Leon Zong, Dylan D. Ray, Lei Lei, Daniel R. Schrider, Franz Baumdicker, Sara Mathieson

## Abstract

Machine learning approaches have become a powerful alternative to traditional methods in population genetics. Convolutional neural networks (CNNs) in particular have been successful in inferring natural selection, recombination rate estimation, introgression, dispersal distances, and effective population size changes. One limitation of CNNs and other deep learning methods is that they can be difficult to interpret. When they have been shown to be as or more successful than summary-statistic-based methods, what are they learning? Here we investigate CNNs from two different methods: the **pg-gan** discriminator for identifying real vs. simulated data, and two networks trained to detect selective sweeps. We first compute correlations between learned network features and traditional summary statistics, then assess whether summary statistics can be predicted from the learned features. To understand the learned features, we compute feature importance through SHAP values and feature groupings through dimensionality reduction. Finally we use decision trees and random forests to build an interpretable “model-of-the-model”. Our results reveal that some CNN architectures can implicitly compute summary statistics such as pairwise heterozygosity, while statistics such as the site frequency spectrum are less similar to the network’s learned features. We find that long-range linkage disequilibrium is readily approximated by the networks and may be more efficiently computed by CNNs than traditional methods (which are quadratic in the number of sites). Overall, this work contributes to the interpretability of deep learning methods in population genetics by clarifying the relationships between model architecture, learned network features, established summary statistics, and predicted evolutionary parameters.

## 1 Introduction

In population genetics, we seek to understand the evolutionary forces shaping genetic variation over time, such as mutation, recombination, natural selection, genetic drift, migration, and population size changes. To study these processes, we often rely on computational tools and statistical methods to make inferences about evolutionary histories from real genetic data. Often this is done by simulating datasets for testing hypotheses, selecting among competing evolutionary models, or inferring model parameters. Recently there has been a shift away from traditional statistical methods (e.g. Approximate Bayesian Computation [8, 9, 17]) toward supervised machine learning (ML) [68]. Early ML methods included support vector machines [62] and random forest models [66, 74]. A next wave of methods included artificial neural networks with summary statistic vectors as the input [14, 41, 46, 52, 57, 58, 70]. Naturally, there is a range of correlated summary statistics, with varying effectiveness for parameter estimation and population genetic tasks such as neutrality tests [2]. Neural networks were able to handle redundancy and make optimal use of information across various summary statistics, but these statistics still needed to be chosen and computed.

Convolutional neural networks (CNNs) directly taking genotype/haplotype matrices as their input were a third generation of models, successfully tackling a range of inference problems including natural selection, recombination rate estimation, introgression, dispersal inference, and effective population size changes [4–6, 10, 15, 16, 23, 28, 36, 64, 73, 80, 83]. Part of the strength of these CNNs is that, because they can handle raw genotype matrices, they need not rely on a predefined set of summary statistics, and thus can be applied to tasks for which appropriate statistics may be lacking [23]. Moreover, the use of predefined statistics may cause valuable information to be thrown out, as CNNs operating on raw data have been shown to outperform summary statistic-based methods on various tasks [23, 36]. However, there is a major drawback to these CNN models: because their inference procedure is not based on known summary statistics, understanding what a model has learned can be difficult [45].

### 1.1 Background on training via simulation, and CNNs from generative adversarial models

Because it is not possible to obtain large training data sets where the precise evolutionary history of real populations is known precisely, machine learning methods for evolution are generally trained on simulated data (but see [65]). Traditionally, simulated population-level data relied on approaches such as coalescent simulation [43], forward-time [55], and resampling [19, 89]. The majority of these methods produce simulated data via explicit evolutionary parameters such as effective population sizes and rates of mutation, recombination, migration, etc. These simulation approaches can be very effective, and can be performed via flexible software tools (e.g. `msms` [21], `simuPOP` [55], `msprime` [37, 38], `SLiM` [30–32]), but they often require in-depth knowledge of the underlying biological processes and extensive parameter tuning. In the context of training machine learning algorithms, or parameterizing other model-based inference methods, severe model misspecification can impair inferential accuracy [66]. However, neural networks themselves can address this problem [50]. For example, Generative Adversarial Networks (GANs) [27] provide a more automatic way to tailor data simulation. GANs, composed of a generator and a discriminator, can learn directly from real population data and generate synthetic datasets that closely mimic the patterns of genetic variation observed in natural populations. GANs are therefore able to capture complex patterns without the need for extensive theoretical assumptions and parameter selection. This capability offers a flexible and efficient approach to simulating population data. Currently in population genetics there are two main types of GANs: those where a generative neural network attempts to create data that is drawn from the same underlying distribution as the training data (i.e. no parameterized evolutionary model) [10, 76–78, 91, 92] and those where the generator includes an evolutionary simulator whose parameters are tuned during the GAN training process [20, 29, 61, 86] (see [90] for a review of generative models in evolutionary biology).

In terms of GANs for demographic inference, `pg-gan` [86] attempts to find evolutionary model parameters that generate simulated data that confuses a CNN discriminator that emits a predicted probability that a given genotype matrix is real. At the end of training, the optimal generator parameters can then be used to simulate data mirroring that of the focal species or population. These GANs can thus be used to elucidate the demographic history of the study population, or to facilitate downstream inference. In a followup study introducing `disc-pg-gan` [61], trained discriminators were fine-tuned to distinguish selected from neutral regions, building on the intuition that regions predicted with high confidence by the discriminator to belong to the “real” class rather than the “simulated” class might be subject to some type of non-neutral evolutionary forces. In the `disc-pg-gan` study, correlations were computed between the values of the last hidden layer of the network and classical summary statistics of the genotype matrix as a way to interpret what the discriminator learned. While machine learning methods aim to automatically identify suitable features for a given task, theoretical results for established summary statistics can also guide machine learning methods [13], which is why we chose to analyze summary statistics. From these correlations, it is clear that some features of the model have a strong positive or negative correlation with statistics. However, some features exhibit almost no correlation with any of the statistical data. This observation motivated our current work, with the goal of understanding what these nodes represent, whether they capture novel statistical features, and how the model achieves its ability to differentiate real from simulated data.

### 1.2 Interpretability for population genetic neural networks

In this study we focus on understanding deep learning models specialized for population genetic tasks, specifically CNN [24] and graph convolutional network (GCNs [44]) architectures. In a generic neural network model, the input (genomic data in the form of a matrix) is passed through one or more hidden layers which ultimately lead to an output layer with the final prediction. A CNN model is a specialized neural network that is particularly effective for analyzing grid-like data, such as images and matrices. CNNs use filters to compute an element-wise dot product over small patches of the input. These filters have learnable weights which are modified during training to extract features of the input and feed them into fully-connected layers to perform classification. The goal of our study is to better understand the information extracted by population genetics CNNs. Interpretability for many biological applications has lagged behind other fields such as image recognition [3, 71, 93] and natural language generation [18, 72]. There are notable exceptions in population genetics where interpretability is analyzed [15, 28, 53, 81], though none of these methods directly investigate the learned features of neural networks and their relationship to interpretable summary statistics.

First we analyze the retrained CNN discriminators of GAN models, focusing on their role in distinguishing between real and simulated population genetic data. We identified which types of population genetic information (as measured by summary statistics) are frequently captured by the CNN’s features, even prior to training, implying that the design of at least some CNN architectures makes them inherently capable of estimating values that are highly correlated with foundational population genetic summary statistics. Indeed we obtain similar findings when examining two different neural networks trained to detect selective sweeps—the GCN from [88] that directly examines the inferred ancestral recombination graph [11, 47, 54] and the ResNet CNN architecture [33] used in the same study. Finally, we more directly examine what information may be influencing a neural network’s decisions by modeling its behavior with decision trees and random forests. Our code for interpreting the GAN-style CNNs is available open-source at https://github.com/mathiesonlab/ss-interpret.

## 2 Methods

### 2.1 CNN Models Studied

In this work we analyze two families of CNNs. The first are the discriminators of trained GAN models. These `pg-gan` CNNs have the same architecture as [61, 86], but are optimized for stability (see Table 1). The inputs to the model are genomic regions of length 50kb, with the middle 36 SNPs extracted. Only biallelic SNPs are included, with the major allele encoded as 0 and the minor allele encoded as 1. Inter-SNP distances (rescaled to fall between 0 and 1) are included as a second channel of the input data, duplicated for each haplotype. To create a family of CNNs, we repeat training 20 times per population, for the populations CEU, CHB, and YRI from the 1000 Genomes Project [1]. Thus, there are 60 discriminators in total, with a small subset that we discard due to predicting the same value for all inputs.

**Table 1:** CNN architecture for both families of CNNs: those trained through a GAN process (pg-gan) and those trained directly on the task of distinguishing real from simulated data. Dropout is applied during training between the final dense layers. The input to these networks is a haplotype alignment with 32 SNPs from a 50kb region. The number of haplotypes varies by population: CEU=198, CHB=206, and YRI=216.

| Layer | Param # |
| --- | --- |
| Conv2D (32, (1,5)) | 352 |
| MaxPooling2D (1,2) | 0 |
| Conv2D (64, (1,5)) | 10,304 |
| MaxPooling2D (1,2) | 0 |
| ReduceSum | 0 |
| Flatten | 0 |
| Dense (64) | 24,640 |
| Dense (64) | 4,160 |
| Dense (1) | 65 |
| **Total** | **39,521** |

To eliminate the potential confounding effects of GAN training (where there is not one loss function, but the generator and discriminator are alternately trying to maximize their own loss functions), we retrained discriminator architectures on the original binary classification task of trying to identify simulated data (scores closer to 0) vs. real data (scores closer to 1). Through this process we created the second family of CNNs (see Table 1). For these retrained discriminators, we used equal amounts of simulated data from each successfully trained GAN above. This led to a set of simulations with somewhat diverse evolutionary parameters, each corresponding to the final parameters estimated by one of the 20 GANs, but all matching (to various degrees) the real data. We repeated the procedure for each of the three populations (CEU, CHB, and YRI), for a total of 60 CNNs in the second family. One CNN is discarded which predicts the same value for every input.

In addition, we examined two pretrained networks from [88] that were used to detect selective sweeps by classifying regions according to five categories: a hard selective sweep with the selected site in the center of the genomic window, hard sweep occurring in the genomic window but not near the center (i.e. outside of the center of 11 sub-windows making up the window), a soft sweep on a previously segregating variant [35] in the center of the window, a soft sweep not near the center, and no sweep. One network is essentially a single channel ResNet [33], a 2D CNN originally intended to classify images, and the other is a custom designed graph convolutional neural network (GCN) that operates on inferred marginal tree sequences (in this case from Relate [75]) along with tree summary statistics computed on these inferred genealogies. The architectures of these networks are described in detail in [88] and summarized in Table 2 and Figure 3 from [33]. Of particular relevance here are the numbers of features in the last hidden layers of the networks. In the case of the CNN, we chose a dimensionality of 64 for the final layer and for the GCN the dimensionality was 256 (before dimensionality reduction via PCA).

**Table 2:** GCN architecture from Whitehouse et. al. [88].

| Layer | Type | Input Shape(s) | Output Shape | Activation |
| --- | --- | --- | --- | --- |
| Node embedding | linear | $(B, T \times N, F)$ | $(B, T \times N, 26)$ | none |
| Skip | concatenation | $(B, T \times N, F),$<br>$(B, T \times N, 26)$ | $(B, T \times N, 26 + F)$ | none |
| Graph convolutions (x6) | GATv2Conv | $(2, E),$<br>$(B, T \times N, 26 + F)$ | $(B, T \times N, 26 + F)$ | ReLU |
| Skip | concatenation | $(B, T \times N, F),$<br>$(B, T \times N, 26 + F)$ | $(B, T \times N, 26 + 2F)$ | none |
| Reshape | | $(B, T \times N, 26 + 2F)$ | $(B, T, N, 26 + 2F)$ | none |
| RNN | GRU (1-layer, unidirectional) | $(B, T, N, 26 + 2F)$ | $(B, T, 256)$ | none |
| Tree-level statistics | concatenation | $(B, T, 12),$<br>$(B, T, 256)$ | $(B, T, 268)$ | none |
| Reshape and pad sequences | | $(B, T, 268)$ | $(B, T', 268)$ | none |
| RNN | GRU (1-layer, unidirectional) | $(B, T', 268)$ | $(B, 256)$ | none |
| Tree-sequence-level (global) stats | concatenation | $(B, 37),$<br>$(B, 256)$ | $(B, 293)$ | none |
| MLP (x2) | fully-connected | $(B, 293)$ | $(B, 256)$ | ReLU |
| output | linear | $(B, 256)$ | $(B, M)$ | Softmax |

$B$ = number of tree sequences per batch (batch size)
$T$ = number of trees in a single sequence
$T'$ = maximum number of trees in a sequence in a given batch, used for padding
$N$ = number of nodes in a single tree
$T \times N$ = matrix of trees × nodes comprising all nodes in a single tree sequence (i.e. the number of nodes across all trees in a single sequence)
$F$ = number of node features calculated for each node in the input tree sequence
$E$ = number of edges in all graphs included in the batch of tree sequences
$M$ = the size of the output (the number of classes for classification, or 1 for regression)
$(..., n)$ = number of features

To explore the behavior of these networks, we simulated selective sweeps via discoal [40] in the same manner as in [67, 88]. Briefly, each example in this dataset consists of 208 haploid individuals under the JPT model of population size change inferred by [7], with all sweep models having a beneficial mutation that reached fixation at time *t* ~ 𝒰(0, 2000) generations ago, with a selection coefficient *s* ~ 𝒰(0.005, 0.1) and for soft sweeps an initial selected frequency *f*_0_ ~ 𝒰(1*/N*, 0.2).

As in [88], the genotype matrix inputs to the ResNet CNN were formatted to be of shape (1, # of individuals, # of polymorphisms) and were padded with zeros to the maximum observed number of polymorphisms in each case. The rows were sorted based on cosine distance using seriation as described in [60] using Google’s OR-tools package [56]; note that the sorting algorithm and similarity metric differ from that used by [23] as we found the seriation approach to perform better on the task of identifying introgressed haplotypes [60]. Each allele for each individual in the input matrix was represented as binary encoding of 0 or 1 (ancestral or derived, respectively).

### 2.2 Summary Statistics

We calculated the following classical population genetics summary statistics, with the goal of understanding if the `pg-gan` CNN models are implicitly or explicitly computing quantities correlated with these statistics.

- **pairwise heterozygosity (***π*): Average number of differences between pairs of sequences.
- **hap:** The number of unique haplotypes in the region.
- **linkage disequilibrium (LD):** LD is calculated by clustering pairs of SNPs based on their inter-SNP distance. We divide these distances into 15 bins (within the 50kb basepair window) and average the correlation *r*^2^ within each one [86].
- **folded site frequency spectrum (folded SFS):** The folded SFS counts mutations with each minor allele frequency, from 1 to *n/*2, where *n* is the number of haplotypes.
- **iHS** integrated haplotype score, as originally developed in [84].
- **Garud statistics** haplotype frequency statistics H1, H2, etc from [25].

For the sweep-detection networks, we examined a different set of summary statistics: those calculated by `diploSHIC` [41], as these statistics were chosen specifically for the problem of sweep detection. These statistics include several of those mentioned above: *π*, average *r*^2^ within a window to summarize LD (referred to as *Z*_*nS*_ [39]), Garud’s *H* statistics, and the number of unique haplotypes (referred to as HapCount by `diploSHIC`). The additional statistics included in `diploSHIC`’s set are:

- **Tajima’s** *D*: A neutrality test that measures the degree of deficit or excess of intermediate-frequency alleles [79].
- *θ*_*H*_ **and Fay and Wu’s** *H*: both from [22].
- **maxFDA:** The maximum derived allele frequency found in a window, which is expected to be elevated near hard selective sweeps [48].
- **Kim and Nielsen’s** *ω*: *ω* captures the excess of LD on both flanks of a hard sweep, combined with the comparative lack of LD between pairs of mutations on either side of the sweep [42].
- **distVar, distSkew, and distKurt:** These statistics examine the distribution of the fraction of differences between all pairs of haplotypes in a sample. *π* is the mean of this distribution, and these three statistics are the variance, skewness, and kurtosis, which are altered by sweeps [41].

Because `diploSHIC` is designed to detect the characteristic spatial skews in diversity patterns produced by a sweep, it measures these statistics in a user-specified number adjacent subwindows spanning a larger window (which here corresponds to our entire simulated region). We followed that convention here when examining the ResNet and GCN networks which were both trained to detect sweeps, used `diploSHIC` to calculate these summary statistics in 11 subwindows (the software’s default value) spanning the simulated region.

### 2.3 Understanding CNN learned features

The outputs of the last hidden layer of a CNN represent potentially sophisticated *learned features* of the data. To understand these learned features and their contribution to the final output, we extract and save the last hidden layer of the model. Based on this last hidden layer, we calculated feature contributions using a variety of existing explanation approaches.

#### 2.3.1 Correlation between CNN features and summary statistics

To test a potential relationship between CNN learned features and summary statistics, we first compute Pearson correlations between the outputs of nodes in the last hidden layer (referred to as ‘learned features’) and summary statistics. Additionally, we investigate how well the learned features can predict each summary statistic by performing a linear regressions for each statistic. The regressions are fit using ordinary least squares with the learned features as input.

#### 2.3.2 SHapley Additive exPlanations (SHAP)

SHAP stands for SHapley Additive exPlanations, which is an approach to help quantify the contribution of each input feature or learned feature to predictions of the model. SHAP can provide both global (overall model behavior) and local (individual prediction) explanations. SHAP is based on Shapley value explanation, which is a game theory principle [69], and is usually used to solve how to distribute the gains of each individual in a cooperative game group. Here, the discrimination task is a teamwork game, and the contribution of the each feature is analogous to the gain attributed to each player in the game.

Compared to normal Shapley value explanation, SHAP introduced a new concept called additive feature attribution, which is defined as adding up all the contributions from different features [49]. By representing Shapley value explanations as an additive feature attribution method (linear model), SHAP effectively combines Shapley values and LIME (Local Interpretable Model-agnostic Explanations) together [51] as shown in Equation 1:

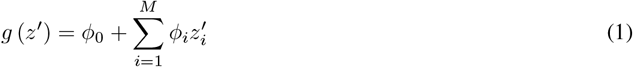

where, *g* is the explanation model, *z*′ ∈ {0, 1}^*M*^ indicates if a feature value is “present” (1) or “absent” (0) [51], *ϕ*_*i*_ ∈ ℝ is the Shapley value for feature *i*, and *M* is the maximum number of features. Every feature is assigned a SHAP value, with a positive value indicating that the presence of the feature increases the prediction output, and a negative SHAP value indicating the feature lowers the prediction. SHAP has become a widely utilized tool in many studies, helping researchers to understand the relationship between input and output in complex ML models. It has been particularly helpful for feature selection and explainability of models [63].

#### 2.3.3 Permutation Explainer

Permutation strategies are commonly used to explain the effect of features on the prediction [12]. The importance of a feature is quantified by observing the increase or decrease in the model’s error after randomly shuffling the values of a feature breaking any potential feature-target relationship. A significant decrease in prediction error indicates that the output relies heavily on the feature, so this feature is important. However, permutation of feature values introduces randomness into the process, reducing reproducibility of computed importance scores. SHAP Permutation Explainer iterates through an entire permutation of each feature in two directions, forward (adding features one by one) and reverse (removing features one by one) to approximate the Shapley values and observe changes in model predictions. This approach provides a global interpretation of the model’s behavior, and can handle larger datasets faster compared to other explainers in SHAP.

In our work, we feed real and simulated data through the CNN models and extract the learned features from the last hidden layer. These learned features are used as the input for calculating SHAP values, with the goal of understanding each of their contributions to the final prediction. SHAP values were computed and visualized using a beeswarm summary plot as implemented in the SHAP python package [49], providing an overview of the importance and impact of each learned feature.

### 2.4 “Model-of-the-model” approach

Moving beyond the last hidden layer, one way to understand a complex neural network is to model its behavior using a simpler, more interpretable model. In [59] the authors created a support vector machine (SVM) method to predict the outcomes of chemical reactions. To understand the SVM, they then trained a C4.5 decision tree to predict the *binary output of the SVM*. In this way they created a simpler model designed to produce outcomes similar to the more complicated SVM. Similar approaches have since been developed for other applications [85, 94].

Decision trees are very interpretable, as they can help transfer complex patterns into simpler, human-understandable decision paths. Thus we followed this approach to create a decision tree model that uses interpretable summary statistics as the features, and a “label” of the CNN’s prediction. In this way we train a more interpretable model to approximate what the CNN has learned, including any prediction errors it has made. Note that this differs from previous approaches in that we are not feeding in the exact same data (i.e. haplotype matrices) but summary statistics computed directly from these matrices.

We additionally train random forest models on the same task. As an ensemble learning method, random forest reduce over-fitting by averaging across multiple trained decision trees. We examine the importance of input features for these random forests in order to reduce one-off effects from examining single decision trees.

## 3 Results

### 3.1 Evaluation of trained CNNs

In interpretability research, if we want to understand what a model has learned, it is important to first assess if the model has learned anything. To that end, we sought to assess the performance of `pg-gan` discriminators across successful `pg-gan` generators, which should all nominally be creating simulations that match the real data very well. We thus ran data from each generator through each discriminator (after eliminating the seeds where the discriminator predicted the same output for all inputs). The results are shown in Figure 1. We note that typically the accuracies are not high, but there is high variance. For example, seed 3 for YRI is more accurate on the real data than the simulations, indicating a propensity to emit higher values (i.e. higher estimated probabilities that examples are real rather than simulated). In contrast, seeds 7 and 12 predict lower values, leading to higher accuracies on the simulations but lower accuracies on the real data.

**Figure 1.**
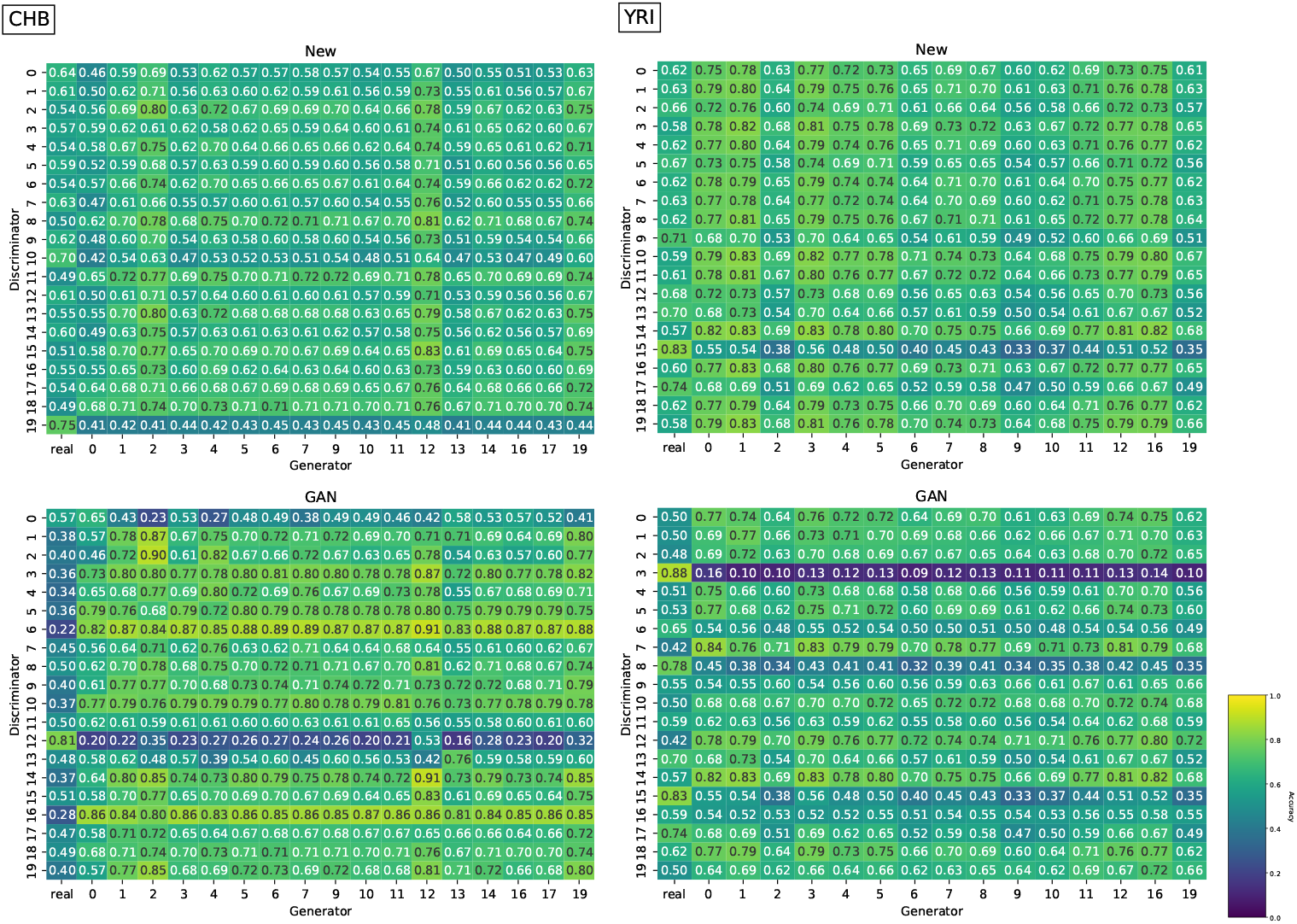
Accuracies of newly trained CNNs (top) and `pg-gan` discriminator CNNs (bottom) for CHB and YRI (see Figure S1 for CEU). Results are separated into real (column 1 of each heatmap) and generated data, and further separated into simulated data from distinct pg-gan generators. Generators are filtered to only those which did not collapse during GAN training. Discriminators are filtered out which predicted the same value for every input.

The discriminators retrained for the original task (distinguishing real from simulated data) had less variance and overall were able to achieve slightly higher accuracy than the original `pg-gan` discriminators, as shown in Figure **??** and S1, and averaged in Table 3. For example, for CEU the accuracy increased from 0.549 to 0.611. (See Figure S2 for a distribution of the raw prediction values separated by real vs. simulated data). However, these are not particularly high accuracies, indicating that either the simulations are very close to the real data, or the differences are too subtle for this particular CNN architecture and training regime to identify. Nonetheless, the fact that these discriminators obtained better than random accuracy on this difficult task implies that they were able to extract features with valuable evolutionary information. Note that we do not evaluate the performance of the ResNet CNN and the GCN trained to detect sweeps, as these were previously shown to perform well on this task [88], and we can therefore assume that these networks are able to extract informative features.

**Table 3:**
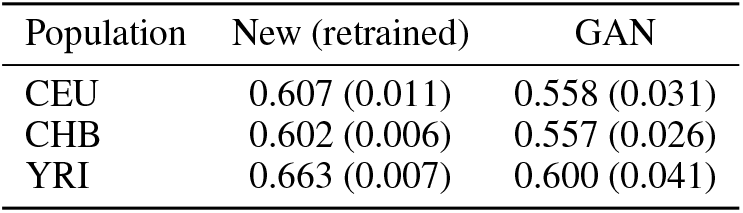
Averaged accuracies of newly trained CNNs (left) and pg-gan CNNs (right) across populations. Results are shown as “mean (SD)” for the task of distinguishing real from simulated data. Overall the newly trained CNNs are not a substantial improvement over the `pg-gan` CNNs, indicating that either the simulations match the real data very well, or this architecture cannot pick up on subtle differences.

| Population | New (retrained) | GAN |
| --- | --- | --- |
| CEU | 0.607 (0.011) | 0.558 (0.031) |
| CHB | 0.602 (0.006) | 0.557 (0.026) |
| YRI | 0.663 (0.007) | 0.600 (0.041) |

### 3.2 Correlations between CNN learned features and summary statistics

To investigate the general relationship between the model’s learned features (here calculated as last hidden layer outputs) and traditional summary statistics, we computed correlation heatmaps (Figure 2). Correlations between learned features and summary statistics were computed for a combined dataset of real and simulated data. We first computed correlations for trained CNNs, as shown for seed 0 in Figure 2, left. We observe strong correlations of some hidden features with the number of singletons (`SFS_1`), short-range LD, *π*, and Garud haplotype statistics [25]. To determined whether these correlations emerge during training, we also computed correlations for a non-trained CNN, i.e. a network of the same architecture but with randomly initialized weights that has never seen any real or simulated training data. In this case (Figure 2, right) we still observe some strong correlations, particularly with *π*. We believe this indicates some implicit capabilities of the *architecture* to compute features that are highly correlated with pairwise heterozygosity, perhaps through synthesizing the output of the 1D convolutional filters (which span one haplotype at a time) through a permutation-invariant function. Indeed, we note that in biallelic data like the data examined here, *π* per site is determined strictly by the major/minor allele frequency at a site. This is equivalent to the sum of the pixel values in a column of the image representation of a haplotype matrix. While the `disc-pg-gan` architecture may not be able to calculate this directly (because convolutions are performed within haplotypes before applying a permutation-invariant function that combines information across haplotypes), it is reasonable to assume that it would readily compute information that is highly correlated with, for example, the number of bright pixels within a small window of sites.

**Figure 2.**
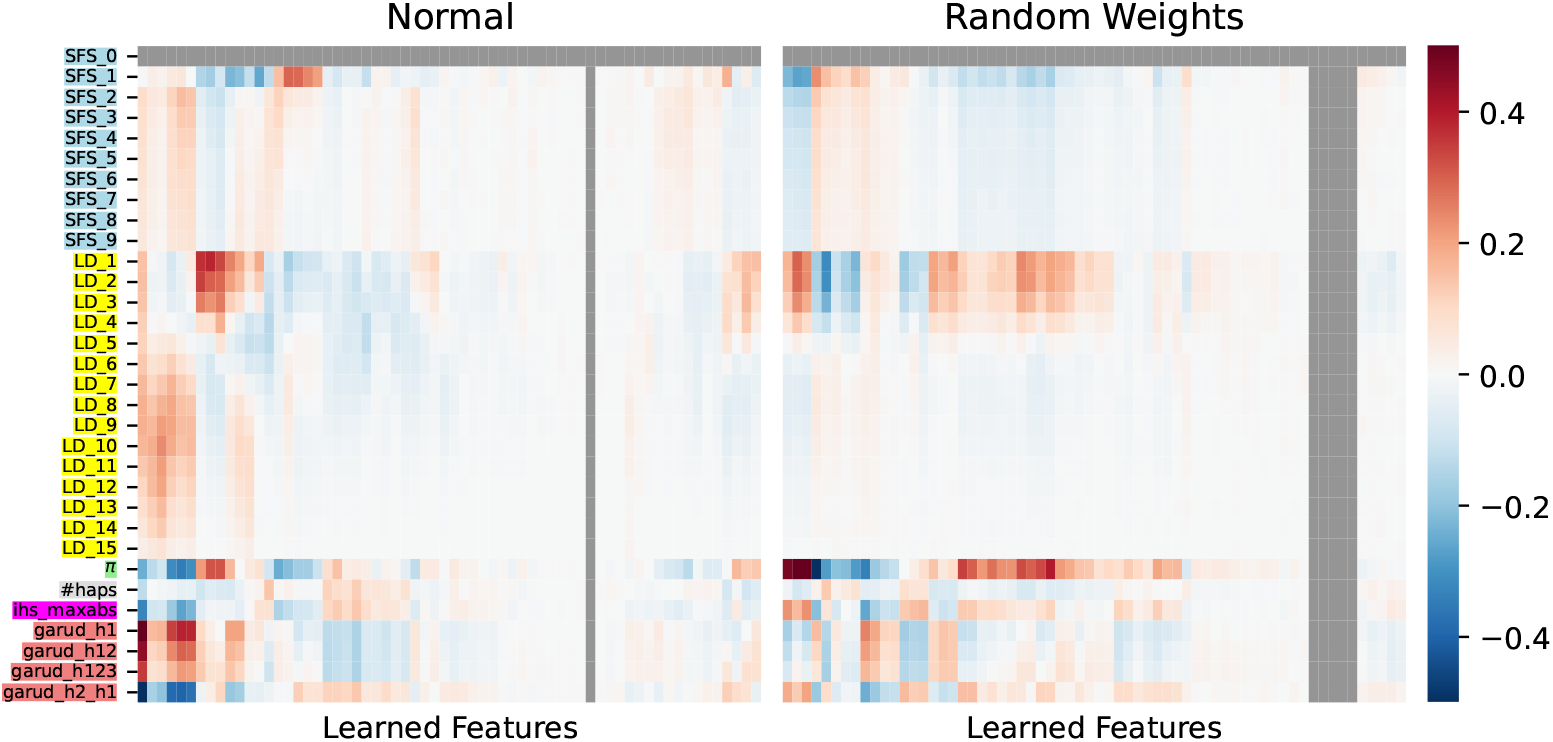
Correlation between last hidden layer values and summary statistics for a trained CNN (left) vs. a randomly initialized CNN (right), seed 0. The *x*-axis shows the 64 features of the last hidden layer, clustered by similarity of their correlation profiles. The *y*-axis shows the summary statistics, restricted to the most informative. Note that we filter non-segregating sites, so `SFS_0` is always 0.

To further understand the differences between a randomly initialized CNN and a trained one, we fit linear regression models to assess the capability of the learned features, as opposed to the randomly initialized features, to predict each summary statistic. The results are shown in Figure 3, with *R*^2^ used to measure agreement between the prediction and true summary statistic value. We observe that a CNN with random weights is easily able to predict *π*, again indicating some innate abilities of the architecture. For statistics where the trained CNN (blue) is better able to predict the statistic than the random CNN (orange), this indicates some learning during training. In our scenario this applies to some of the haplotype frequency statistics (e.g. `garud_h1` and `garud_h2_h1`) as well as the mid-long range LD statistics (e.g. `LD_7` – `LD_11`). One takeaway of this result is that long-range LD is being learned by our models, although it may or may not be useful for the final prediction. If long-range LD is useful for identifying real vs. simulated data, it could indicate a shortcoming of our simulation framework available for the GAN-generator (i.e. unmodeled population structure and non-random mating patterns).

**Figure 3.**
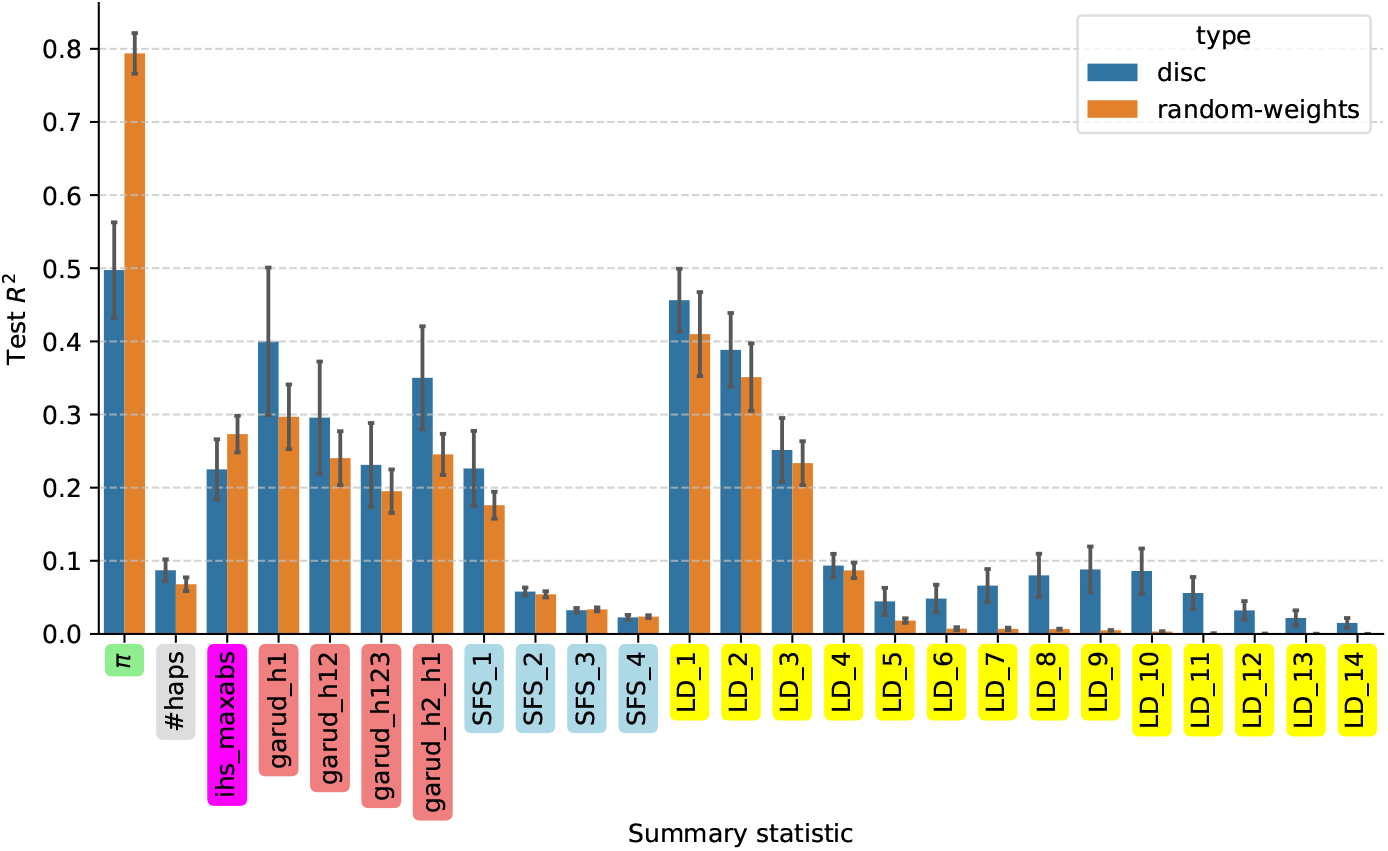
*R*^2^ achieved by modeling the relationship between the last hidden layer values and the summary statistics with a linear regression. Blue bars indicate trained CNNs and orange bars indicated non-trained CNNs (randomly initialized weights). Error bars indicate standard deviation across 5 seeds.

Notably, the agreement of the features with *π* is smaller for of the trained CNN than the un-trained CNN, indicating that although easily achievable an exact estimate of pairwise diversity is not crucial for the prediction. In addition to *π*, short-range LD is implicitly computed by an untrained CNN. This could be explained by the range of the convolutional filters. Considering both convolutional layers, the filters can access SNPs up to 16 indices apart. On average, this is 3524 bases apart, which encompasses LD bins 1-3. Our `LD_4` bin corresponds to SNPs between [4286, 5714] bases apart, which is beyond the range of the convolutional filters such that random weights less likely generate correlating features. This could explain the LD pattern we observe, with implicit computation of short-range LD and learned computation of mid- and long-range LD.

We ran the same analysis for the CNN and GCN model from [88] that were trained to detect selective sweeps by distinguishing between five classes, four corresponding to hard and soft sweeps either at or far from the center of the simulated region, and one neutral class (Methods). We simulated 1000 replicates for each class (or 5000 in total) and computed summary statistics using `diploSHIC` (15 statistics over 11 windows of the simulated chromosome). Again, we observed substantial correlation between the networks’ features (both before and after training) and population genetic summary statistics (Figures S3 and S4). Overall there are larger differences between the trained and untrained networks in terms of their ability to predict summary statistics (Figures 4 and 5). Statistics such as *π*, Watterson’s *θ* (*θ*_*w*_), or *θ*_*H*_ can be reconstructed with decent accuracy (*R*^2^ *>* 0.5 for each, and *R*^2^ *>* 0.8 in the case of *θ*_*w*_) from features from a randomly initialized network and remain so in a trained network, which again may be expected given that these summaries are correlated with the number of major/minor alleles and/or number of segregating sites in our haplotpye matrix. However, statistics such as the maximum derived allele frequency (maxFDA), the number of distinct haplotypes, and Garud’s *H* statistics are found to be more recoverable after training. In the case of GCN, *π* and *θ*_*w*_ have higher *R*^2^ scores for a random network than after training, and many statistics have higher *R*^2^ values from the untrained GCN than from the untrained ResNet CNN.

**Figure 4.**
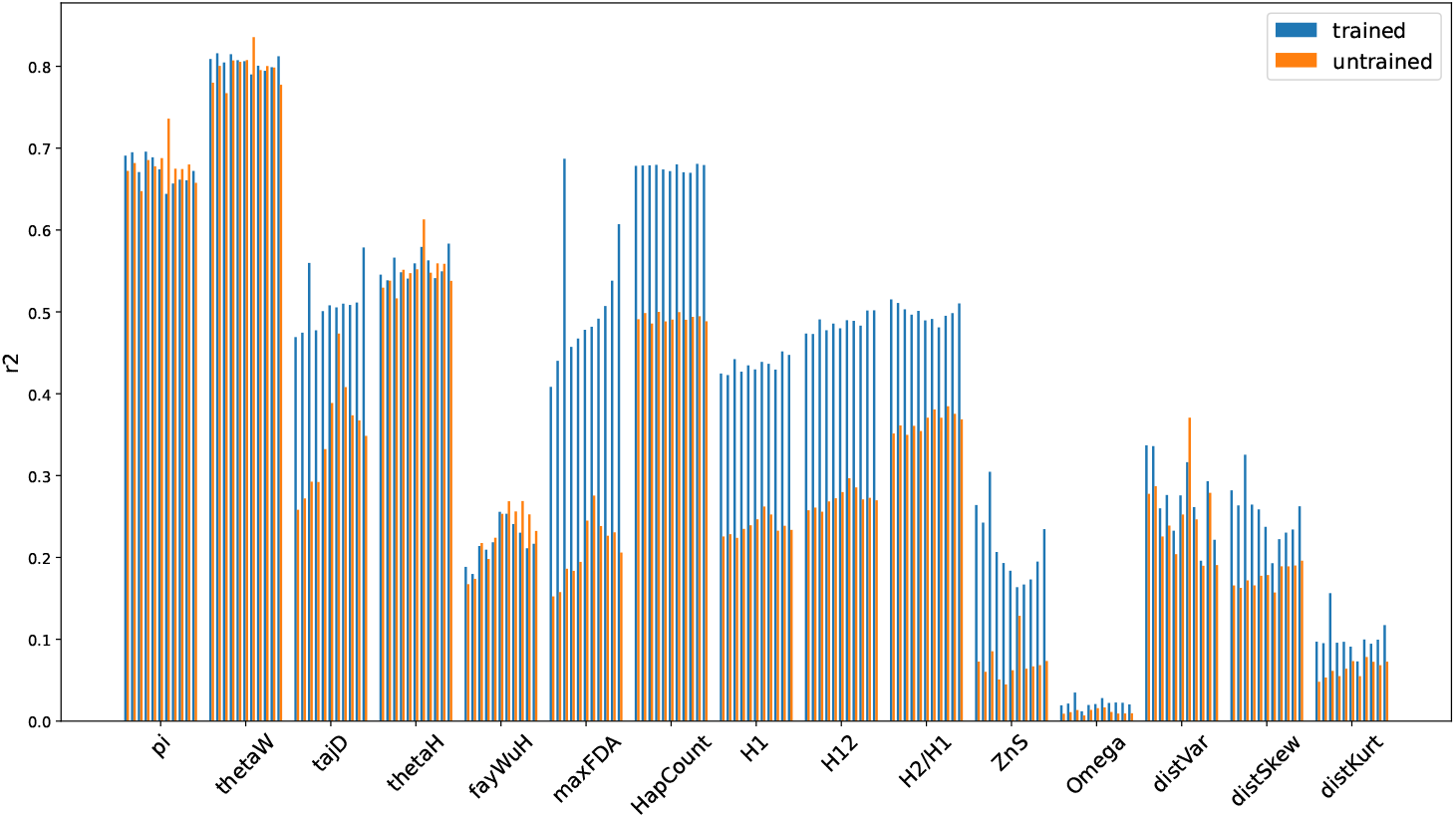
*R*^2^ achieved by modeling the relationship between the transformed last hidden layer values and the summary statistics with a linear regression (for the ResNet sweep-detection CNN from [88]). Blue bars are from a trained CNN and orange bars are from a non-trained CNN (randomly initialized weights).

**Figure 5.**
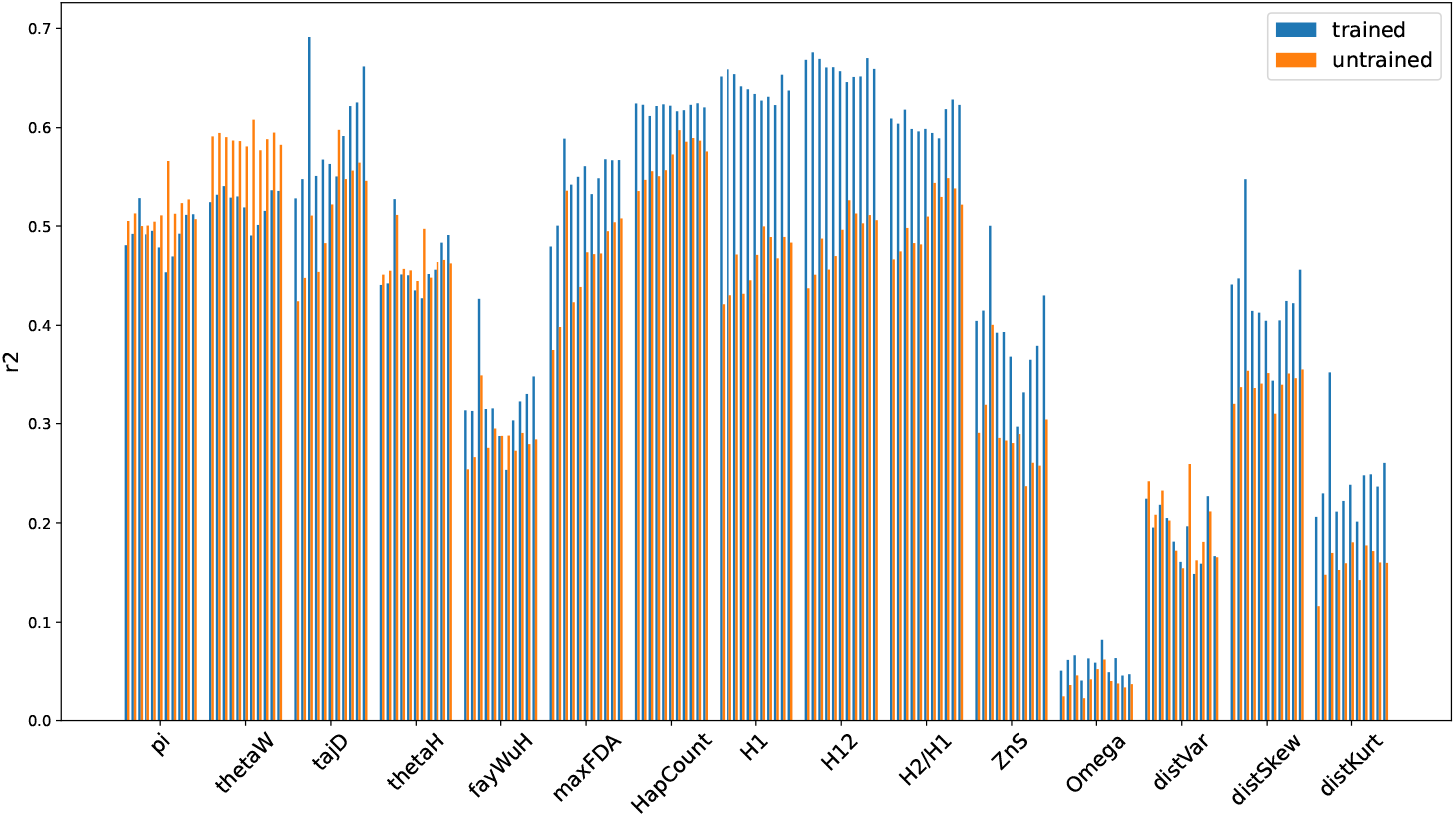
*R*^2^ achieved by modeling the relationship between the transformed last hidden layer values and the summary statistics with a linear regression (for the GCN sweep-detection model from [88]). Blue bars are from a trained GCN and orange bars are from a non-trained GCN (randomly initialized weights).

**Figure 6.**
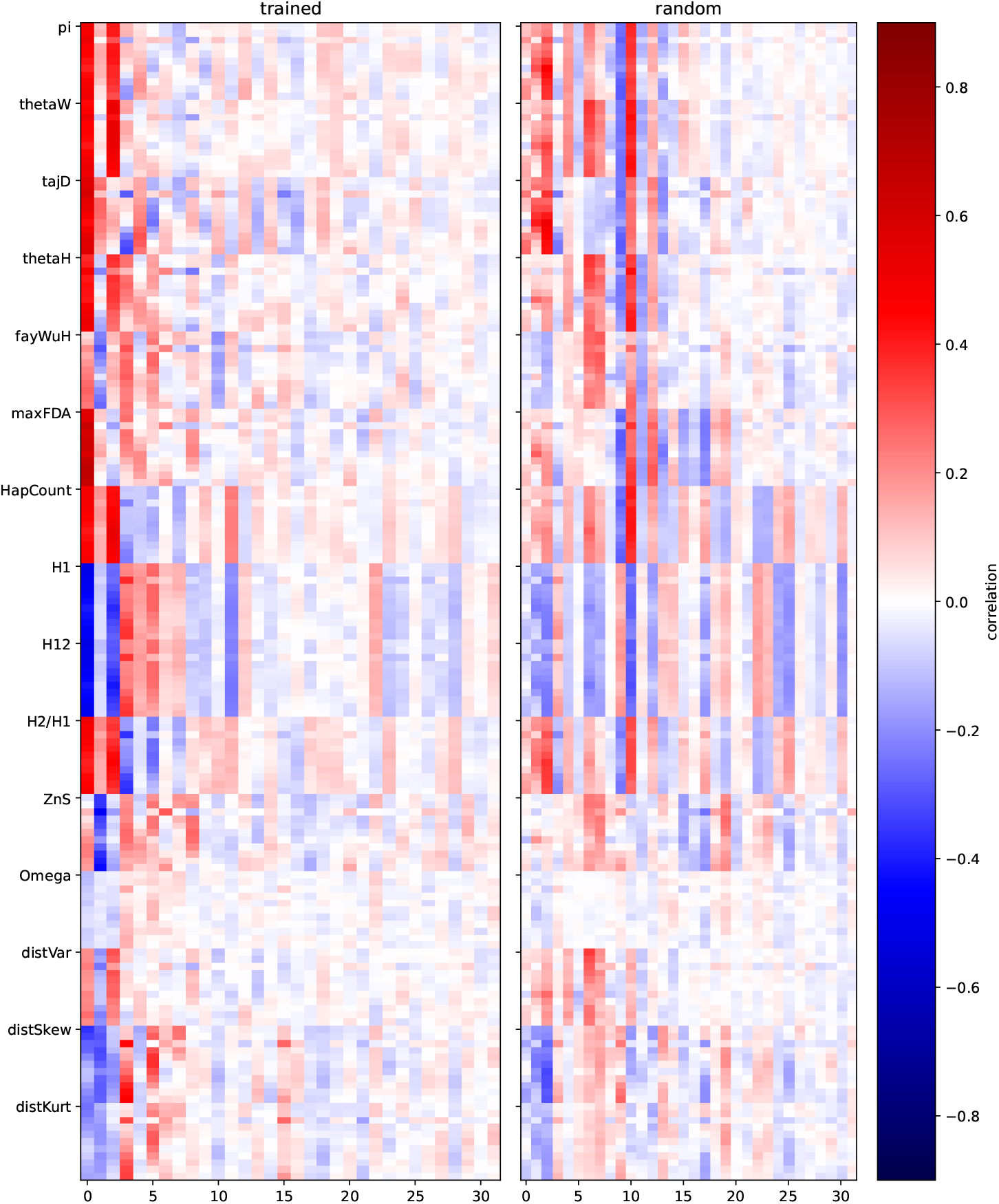
Side by side comparison of correlations of transformed GCN (first 32 principal components) features with `diploShic` summary stats. For a pre-trained network on the left and random weights on the right.

### 3.3 Dimensionality reduction and SHAP analysis of learned features

Based on the correlation heatmaps between the last layer values and summary statistics (Figures 2, S5, and 6), some learned features seem to capture similar information to one another. To see the features and their associated class membership probabilities without this redundancy, we performed a principal component analysis (PCA) transformation to reduce the dimension of the 64 learned features of the `disc-pg-gan` discriminators. We plot the learned features of the combined real and simulated data with prediction values to distinguish potential features that might affect model performance (Figure 7).

**Figure 7.**
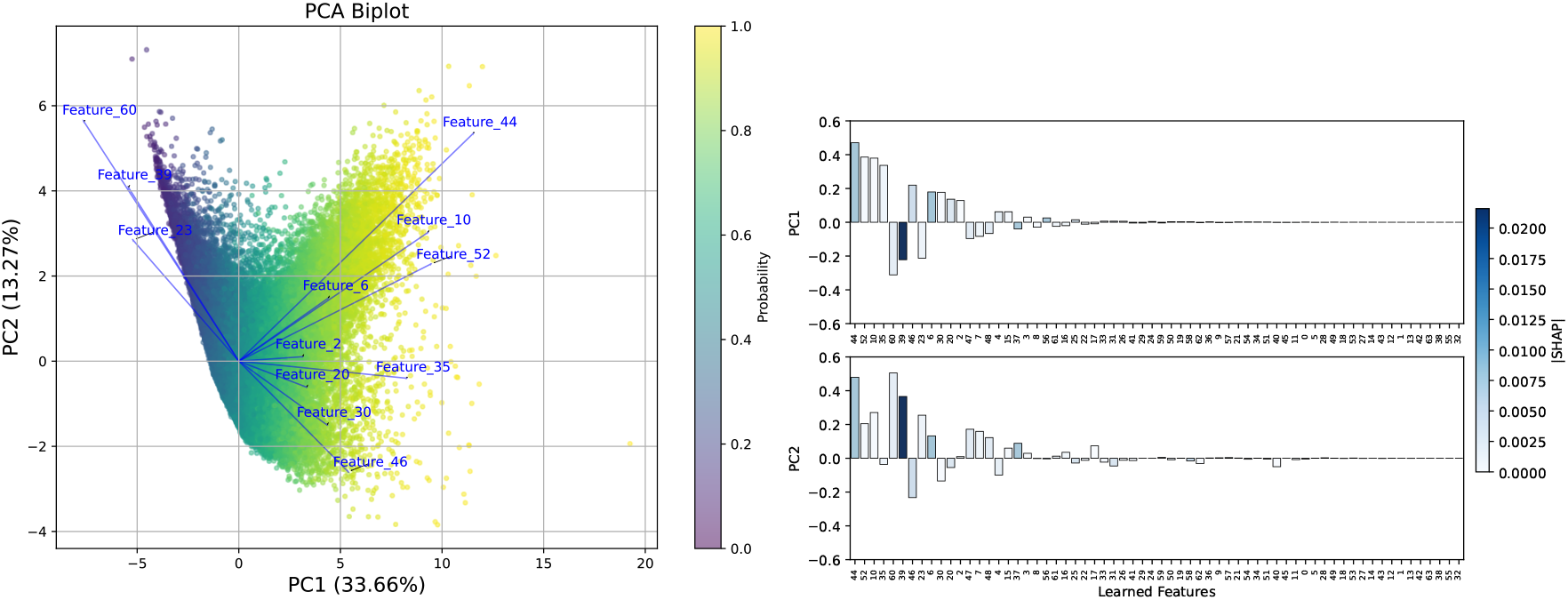
Learned feature visualizations for a CNN trained with CEU data, seed 0. **Left:** PCA visualization for real and simulated last layer values colored by probability. The x-axis represents the first principal component (PC1), explaining 33.66% of the variance, and the y-axis represents the second principal component (PC2), explaining 13.27% of the variance. The points are colored according to the model’s predicted probability, yellow (value close to 1) indicates a high prediction of being real data, and purple (value close to 0) indicates a high prediction of being simulated data. This biplot illustrates the contributions of each learned feature to the first principal component (PC1). Each point represents the last layer values of simulated and real regions, transformed by PCA and rescaled to range from 0 to 1. Arrows show feature contribution vectors (threshold 0.1). **Right:** Bar plot of the learned features contributions to PC1 and PC2. The x-axes show the 64 learned features of the last hidden layer, and the y-axes show their respective contributions to PC1 and PC2. The learned features are ordered by the absolute value of their contributions to PC1 and colored by mean absolute SHAP value. The color gradient is based on mean absolute SHAP value, where deeper blue means higher SHAP values, indicating more importance for the model’s predictions.

These PCA scatter plots demonstrate that the prediction values can be quite distinct based on the provided last layer features. PC1 and PC2 are able to explain a large portion (46.93%) of the variance. The scatter plot exhibits a “check mark” shape, with the head of the check mark corresponding to predicting simulated data (probabilities closer to 0), and the tail corresponding to to predicting real data (probabilities closer to 1). Distinct differences in the distributions of the real and simulated datasets are observed, showing that the values of the last layer in the model capture the variance between these two datasets.

To better understand the PCA results and determine which features have an impact on these two directions respectively, we focused on the first two principal components (PC1 and PC2), which explain 33.66% and 13.27% (respectively) of the variance in the merged dataset. A biplot and a bar plot are generated to visualize the contributions of features to PC1 and PC2 (Figure 7). The arrows in the biplot represent the vectors of each feature, with their directions determined by the contributions to PC1 (X-axis) and PC2 (Y-axis). The length of each arrow reflects the amount of the feature’s contribution. Figure 7 indicates that features on the left (e.g. indices 23, 39, and 60) are related to a lower probability prediction (i.e. higher values of these features are indicative of simulated data). Features on the right (e.g. indices 10, 44, and 52) are indicative of a higher probability prediction (i.e. higher values of these features are indicative of real data). Results are shown for CEU seed 0, so different learned features would appear for different CNNs, but the general patterns hold.

On the right of Figure 7 we show the contributions of learned features to PC1 and PCs, colored by their SHAP values. (See Figure S6 for deep SHAP and permutation SHAP results.) We note that generally the nodes with lower correlation with summary statistics statistics have lower SHAP values, suggesting that they have lower influences on the model’s output and likely do not capture meaningful information.

The features extracted from the GCN and ResNet CNNs for sweep detection (trained and randomly initialized weights) were also highly self-correlated, so we again performed PCA on these data. We see that correlations are generally lower for the random weights than those for the trained network, especially for the higher principal components. This again supports the notion that, while these neural network architectures extract information that is correlated with commonly used summary statistics even without training, training causes the network to learn features that are even more strongly correlated with these statistics.

### 3.4 Random forest models

Following the “model-of-the-model” approach outlined in the Methods section, we train a random forest regression model to predict the probability outputs of the *original* model, given summary statistics for each region as input. The results are shown for each population in the bottom row of Figure 8. As a baseline, we also train a random forest classifier on the original task (binary prediction of real vs. simulated data), again using summary statistics as the input. This experiment does not shed light on what a neural network model is learning, but instead identifies which summary statistics would a summary-statistic-based model find most useful for this task. The accuracy of these models are shown for each population in the top row of Figure 8, and can be compared to Table 3.

**Figure 8.**
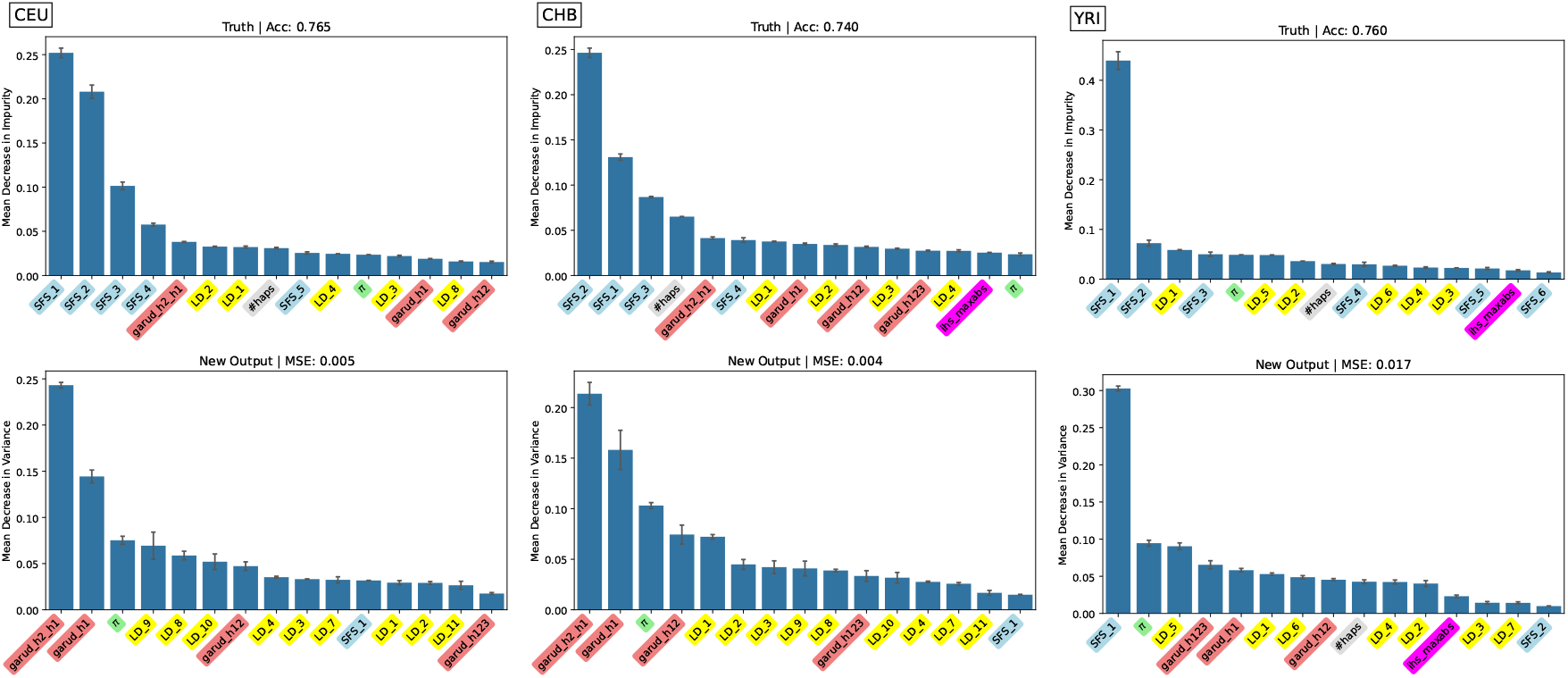
Feature importance of random forest models trained to classify real versus simulated data (top row) and trained to regress trained CNN output with the same task (bottom row). Feature importance is mean decrease in impurity for classification and mean decrease in variance for regression. All random forests use summary statistics as input and importance is shown for each population.

The random forest models on the original classification task had fair accuracy (0.74–0.765), although notably higher than the CNN models (0.602–0.663, see Table 3). This is additional evidence that the simulations match the real data quite well overall. The discrepancy between the random forest models and CNN models can be explained in part by the CNN’s difficulty with computing the site frequency spectrum (SFS), which is provided to the random forest. As shown in the feature importance (Figure 8, top row), SFS values for rare variants were particularly important in distinguishing real from simulated data, possibly due to sequencing errors, rare variant filtering, or inference error in the exponential growth parameter. The CNN architectures analyzed here might have a particularly difficult time estimating rare variant counts due to smoothing and combining of convolutional filters which span multiple SNPs at a time.

For random forest models trained to predict the CNN output, (Figure 8, bottom row), we find that haplotype statistics and *π* are very important, as well as LD statistics at various SNP distances. This analysis sheds light into how the CNNs are making their predictions.

Finally, to construct an interpretable model that seeks to mirror the behavior of the CNN, we restrict our analysis to a single decision tree and truncate to depth 3 (Figure 9). The bottom tree mirrors the results of Figure 8 (bottom row) with haplotype statistics and mid-range LD statistics enabling the decision tree to predict the CNN output. In particular, if mid-long range LD is higher than expected (`LD_8` – `LD_11` here) then data is predicted as more “real”. This could indicate unmodeled population structure, which would often elevate long-range LD. This result is also mirrored in Figure 2 (where we see mid-long range LD correlated with some learned features) and Figure 3 (where learned features can predict `LD_7` – `LD_11` much better than a randomly initialized CNN). The Garud haplotype statistics have a similar pattern, where they are correlated with learned features (Figure 2), can be predicted from the trained network (`h1` and `h2_h1` in particular, Figure 3), are very important for predicting the CNN output (Figure 8, bottom row), and appear in the bottom decision tree (Figure 9). Again, for comparison we also trained a decision tree to distinguish between real and simulated examples. This is shown in the top tree of 9, which mirrors the results of Figure 8 (top row) with rare variant frequencies being the most distinguishing summary statistics between real and simulated data.

**Figure 9.**
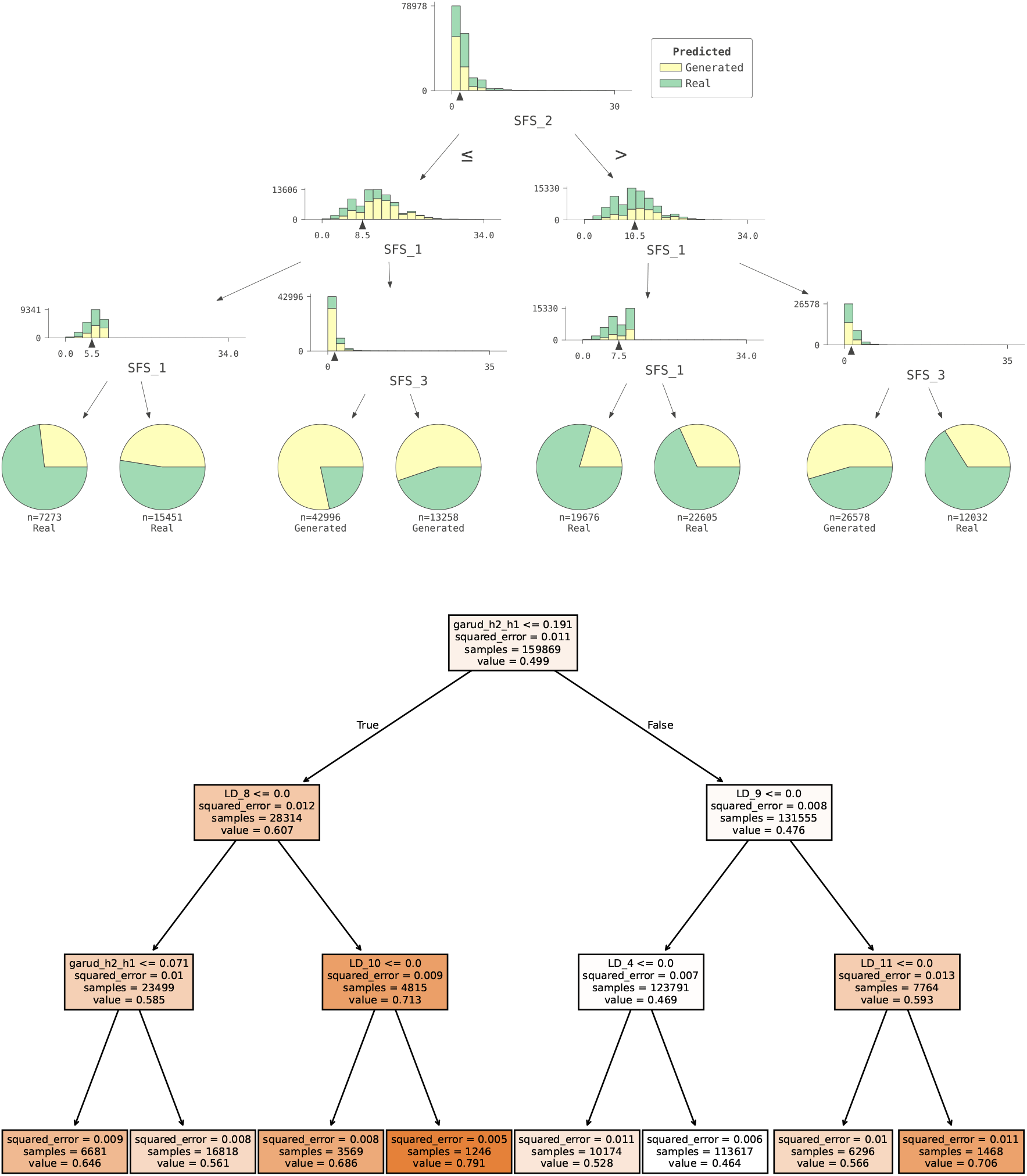
Truncated example decision trees (depth 3). **Top:** modeling the ground truth task of real vs. simulated data (binary classification task) with summary statistics as input. Green indicates real data predictions and yellow indicates simulated data predictions. **Bottom:** modeling CNN output predictions (regression task) with summary statistics as input. Darker orange colors represent real data predictions and lighter orange colors represent simulated data predictions.

## 4 Discussion

Understanding the internal mechanisms of machine learning models is important for trusting their prediction results and characterizing the differences between the different classes under consideration. For example, in the context of `pg-gan`, probing the behavior of the discriminator can reveal features of real data that are well modeled in the simulations. In this study, our goal was to understand the relationship between complex CNN architectures and more interpretable summary statistics. We showed several ways to interpret the learned features of the discriminator that together create a more comprehensive understanding of the CNN predictions. First, the learned features can be interpreted by analyzing their relationship with known summary statistics. Second, PCA can be applied to understand redundancy in the learned features, providing insights into feature contributions. Third, SHAP values can offer global or local explanations of feature importance. Lastly, we used random forest and decision tree models to predict the CNN output as well as the output for the original task, to understand which types of information the network might be using to reach its final prediction.

There are a few other works in population genetics that have analyzed interpretability. Cecil and Sugden [15] also use SHAP values, though directly on the haplotype matrices to see which aspects are most important for selection. Gower et al. [28], Whitehouse and Schrider [87], and Nait Saada et al. [53] all use saliency maps [71], again on the input data. Tran et al. [81] address interpretability most directly – in this study the authors perturb the haplotype matrix input in various ways to assess the impact of the downstream inference. They also find that linkage disequilibrium patterns are important for selection (for some networks) and allele frequency information is more important for some other applications. We distinguish our methods as focusing on the learned features of neural networks and understanding their relationship to traditional summary statistics.

To remove the effects of GAN training on the resulting CNN discriminators, we trained CNNs of the same architecture on the original task for distinguishing real vs. simulated data. We found that the accuracy was not much higher than the GAN-trained CNNs, indicating either this is a difficult task for this architecture, or the simulations closely match the real data. Later experiments with random forest models lend support to the idea that the simulations are relatively difficult to distinguish from real data, as no method achieved anywhere close to 100% accuracy. This is in contrast to some simulation work in phylogenetics, which suggests it is relatively easy to distinguish real from simulated data [82] with neural network methods. For us, it is possible that if methods were retrained to distinguish real data from simulated data generated from only *one* seed (i.e. one set of evolutionary parameters), our accuracy would increase.

Correlation analyses between individual learned features in the last hidden layer and calculated summary statistics showed that the relationships were not particularly strong. The maximal absolute correlation coefficient is not much higher than 0.4, with the exception of a few statistics in the case of the sweep-detection networks. The general lack of strong correlations indicates that the discriminator might learn more complex nonlinear combinations of features that are not captured by summary statistics, or it is possible that the model learned a more dense representation of summary statistics. The learned features with lower correlation with statistics have lower SHAP values, suggesting that they have lower influences on the model’s output and likely do not capture meaningful information. This aligns with PCA findings, where the features with fewer contributions in PC1 have lower SHAP values, and dominant features in PC1 are also highly ranked by SHAP. Contrary to our expectations, we did not find evidence that the learned features with low correlations with classical population genetics statistics could in fact represent “novel summary statistics”. Some of these learned features have essentially zero variance, so it is likely they do not contain any biological meaning. However, those with relatively low but non-zero SHAP values may contain some information. If CNNs are not learning truly novel features, the question of how they obtain better performance than summary statistic-based methods remains open. One possibility is that the features that they are learning, while correlated with traditional summary statistics, contain slightly more discriminatory value than these statistics (i.e. CNNs are learning “improved” rather than “new” statistics). Similar arguments have been outlined for traditional approaches where instead of a fixed summary statistic a class of summary statistic variants is considered for neutrality tests [2].

One interesting takeaway from our work is that a randomly initialized (untrained) CNN can compute quantities that allow for the accurate prediction of *π*. We speculate that the network is not computing *π* by analyzing all pairs of haplotypes, nor is it directly computing allele frequencies at each site and using those to calculate the number of pairwise differences between haplotypes. Instead, the network might be approximating the number of minor alleles within windows through the initial convolutional layers, which can be used to predict *π* non-linearly. If the deeper layers of the network can approximate this quadratic function, then an estimate of *π* can be computed without directly counting allele frequencies or differences between all pairs of haplotypes. In this way the network could predict how many pairs would have differences along the region.

On the flip side, our `pg-gan` architecture is not well-equipped to compute the site frequency spectrum. This could be because fine-scale variation is smoothed out by convolutional filters that span multiple SNPs. However, our random forest model based on summary statistics indicates that the SFS would be a valuable feature to discriminate real and simulated datasets. If the SFS is manually computed, then methods like random forests can take advantage, as rare variation was a distinguishing feature between real and simulated data. Overall the SFS signal for real vs. simulated data could be particularly difficult even if the demography is well inferred due to sequencing errors, variant calling filters, and difficulties of inferring recent exponential growth parameters. In our scenario, a method that, due to its architecture, does not capture this kind of information might actually be preferable. Nevertheless, in general, potentially valuable summary statistics that are easy to calculate but hard to infer for the CNN could be manually fed as another channel into a CNN model.

Since we are analyzing the application of distinguishing real vs. simulated data, features that are particularly helpful for this task can help us understand what aspects of real data are not modeled well by our simulations. Our results indicated that elevated mid-long range LD was indicative of real data, though this varies by population. This result points to unmodeled population structure (non-random mating) which is a key area to work on for future realistic simulations. In [26], elevated linkage disequilibrium was also identified as a prominent genomic feature potentially indicative of natural selection in *Drosophila melanogaster*, although selection probably has not had as pervasive an effect on genetic variation in the human genomic data examined here.

There are many avenues for future work related to the interpretability of complex ML models for population genetics. Our results indicate that CNNs may be computing or approximating traditional summary statistics. We wouldn’t need an alternative to compute a statistic like the SFS since it is fast and simple, but computation costs for summaries of linkage disequilibrium are quadratic in the number of haplotypes, making them challenging to compute. It is possible that future CNN models could be trained to approximate LD statistics, which would significantly improve the runtime. This possibility is supported by the fact that CNNs can accurately infer recombination rates and detect hotspots [16, 23]. Another direction would be to explore other layers of the network, including understanding the convolutional filters themselves. They could be picking up on short-range LD or even motifs for other applications. For example, for neural network architectures like the ResNet sweep detector examined here, initial convolutional layers identifying contrasting pixel values along the y-axis of an image could also be extracting information that is predictive of *π* (e.g. the number of pairwise differences between those pairs of haplotypes that happen to be located adjacent to one another in the input image). Here the applications are real vs. simulated data and neutral vs. positively selected regions, but the approaches presented here could also apply to CNNs for other applications such as recombination, dispersal inference, or other types of selection such as balancing selection, background selection, or adaptive introgression.

Finally, we could consider different summary statistics, including tree- or ARG-based statistics [34, 88], (although the actual trees are part of the input to the GCN, so it would not make sense in this case). It would be particularly interesting if haplotype-based neural networks were approximating ARG-related statistics along the region (i.e. number of remaining lineages over time, etc). The idea that networks are computing novel statistics is still appealing, but our results suggest that these networks are instead primarily relying on features that are at least somewhat similar to existing statistics. However, this does not mean that neural networks cannot help elucidate evolutionary processes. For example, we envision that the model-of-the-model approach used here could reveal novel summary statistic combinations that both illuminate the manner in which different evolutionary phenomena affect patterns of genetic diversity, and enable us to detect and characterize these phenomena with both high accuracy and interpretability.

## Acknowledgments

We would like to acknowledge Henry Britton and Kai Britt who performed initial interpretability experiments. DDR and DRS were supported by the National Institutes of Health (NIH) under award number 2R35GM13826. SM is funded in part by a NIH grant R15HG011528. FB is funded by the Deutsche Forschungsgemeinschaft (DFG, German Research Foundation) under Germany’s Excellence Strategy – EXC number 2064/1 – Project number 390727645, and EXC 2124 – Project number 390838134. FB and HX were supported by the Reinhard Frank-Stiftung. The content is solely the responsibility of the authors and does not necessarily represent the official views of the National Institutes of Health.

## References

[1] 1000 Genomes Project Consortium. 2015. A global reference for human genetic variation. Nature, 526, (7571) 68.

[2] Achaz, G. 2009. Frequency spectrum neutrality tests: one for all and all for one. Genetics, 183, (1) 249–258.

[3] Adebayo, J., Gilmer, J., Muelly, M., Goodfellow, I., Hardt, M., and Kim, B. 2018. Sanity checks for saliency maps. Advances in neural information processing systems, 31,.

[4] Adrion, J. R., Galloway, J. G., and Kern, A. D. 2020. Predicting the landscape of recombination using deep learning. Molecular biology and evolution, 37, (6) 1790–1808.

[5] Arnab, S. P., Campelo dos Santos, A. L., Fumagalli, M., and DeGiorgio, M. 2025. Efficient detection and characterization of targets of natural selection using transfer learning. Molecular Biology and Evolution, 42, (5) msaf094.

[6] Arnab, S. P., Campelo dos Santos, A. L., Fumagalli, M., and DeGiorgio, M. 2025. Semi-supervised detection of natural selection with positive-unlabeled learning. bioRxiv, pages 2025–08.

[7] Auton, A., Abecasis, G., and Altshuler, D. October 2015. A global reference for human genetic variation. Nature, 526, 68–74.

[8] Beaumont, M., Zhang, W., and Balding, D. December 2002. Approximate bayesian computation in population genetics. Genetics Society of America, 162, (4) 2025–2035.

[9] Beaumont, M. A. 2010. Approximate bayesian computation in evolution and ecology. Annual review of ecology, evolution, and systematics, 41, (1) 379–406.

[10] Booker, W. W., Ray, D. D., and Schrider, D. R. 2023. This population does not exist: learning the distribution of evolutionary histories with generative adversarial networks. Genetics, 224, (2) iyad063.

[11] Brandt, D. Y., Huber, C. D., Chiang, C. W., and Ortega-Del Vecchyo, D. 2024. The promise of inferring the past using the ancestral recombination graph. Genome Biology and Evolution, 16, (2) evae005.

[12] Breiman, L. October 2001. Random forests. Machine Learning, 45, (1) 5–32. ISSN 1573-0565. doi: 10.1023/A:1010933404324. URL https://doi.org/10.1023/A:1010933404324.

[13] Burger, K. E., Pfaffelhuber, P., and Baumdicker, F. 2022. Neural networks for self-adjusting mutation rate estimation when the recombination rate is unknown. PLOS Computational Biology, 18, (8) e1010407. doi: 10.1371/journal.pcbi.1010407.

[14] Caldas, I. V., Clark, A. G., and Messer, P. W. 2022. Inference of selective sweep parameters through supervised learning. bioRxiv, pages 2022–07.

[15] Cecil, R. M. and Sugden, L. A. 2023. On convolutional neural networks for selection inference: Revealing the effect of preprocessing on model learning and the capacity to discover novel patterns. PLoS computational biology, 19, (11) e1010979.

[16] Chan, J., Perrone, V., Spence, J., Jenkins, P., Mathieson, S., and Song, Y. February 2018. A likelihood-free inference framework for population genetic data using exchangeable neural networks. Neural Information Processing Systems, pages 8594–8605.

[17] Csilléry, K., Blum, M. G., Gaggiotti, O. E., and François, O. 2010. Approximate bayesian computation (abc) in practice. Trends in ecology & evolution, 25, (7) 410–418.

[18] Dang, Y., Huang, K., Huo, J., Yan, Y., Huang, S., Liu, D., Gao, M., Zhang, J., Qian, C., Wang, K., et al. 2024. Explainable and interpretable multimodal large language models: A comprehensive survey. arXiv preprint arXiv:2412.02104.

[19] Efron, B. Bootstrap methods: another look at the jackknife. In Breakthroughs in statistics: Methodology and distribution, pages 569–593. Springer, 1992.

[20] Eneli, A. A., Siu, P. C., Perez, M. F., Burt, A., Fumagalli, M., and Mathieson, S. 2025. On the use of generative models for evolutionary inference of malaria vectors from genomic data. bioRxiv, pages 2025–06.

[21] Ewing, G. and Hermisson, J. 2010. MSMS: a coalescent simulation program including recombination, demographic structure and selection at a single locus. Bioinformatics, 26, (16) 2064–2065.

[22] Fay, J. C. and Wu, C.-I. July 2000. Hitchhiking under positive darwinian selection. Genetics, 155, 1405–1413.

[23] Flagel, L., Brandvain, Y., and Schrider, D. R. 2019. The unreasonable effectiveness of convolutional neural networks in population genetic inference. Molecular biology and evolution, 36, (2) 220–238.

[24] Fukushima, K. 1980. Neocognitron: A self-organizing neural network model for a mechanism of pattern recognition unaffected by shift in position. Biological cybernetics, 36, (4) 193–202.

[25] Garud, N., Messer, P., Buzbas, E., and Petrov, D. February 2015. Recent selective sweeps in north american drosophila melanogaster show signatures of soft sweeps. PLoS genetics, 11, e1005004.

[26] Garud, N. R. and Petrov, D. A. 2016. Elevated linkage disequilibrium and signatures of soft sweeps are common in drosophila melanogaster. Genetics, 203, (2) 863–880.

[27] Goodfellow, I., Pouget-Abadie, J., Mirza, M., Xu, B., Warde-Farley, D., Ozair, S., Courville, A., and Bengio, Y. 2020. Generative adversarial networks. Communications of the ACM, 63, (11) 139–144.

[28] Gower, G., Picazo, P. I., Fumagalli, M., and Racimo, F. 2021. Detecting adaptive introgression in human evolution using convolutional neural networks. Elife, 10, e64669.

[29] Gower, G., Picazo, P. I., Lindgren, F., and Racimo, F. 2023. Inference of population genetics parameters using discriminator neural networks: an adversarial monte carlo approach. bioRxiv, pages 2023–04.

[30] Gower, G., Pope, N. S., Rodrigues, M. F., Tittes, S., Tran, L. N., Alam, O., Cavassim, M. I. A., Fields, P. D., Haller, B. C., Huang, X., et al. 2025. Accessible, realistic genome simulation with selection using stdpopsim. bioRxiv.

[31] Haller, B. C. and Messer, P. W. 2019. Slim 3: forward genetic simulations beyond the wright–fisher model. Molecular biology and evolution, 36, (3) 632–637.

[32] Haller, B. C., Galloway, J., Kelleher, J., Messer, P. W., and Ralph, P. L. 2019. Tree-sequence recording in slim opens new horizons for forward-time simulation of whole genomes. Molecular ecology resources, 19, (2) 552–566.

[33] He, K., Zhang, X., Ren, S., and Sun, J. Deep residual learning for image recognition. In Proceedings of the IEEE conference on computer vision and pattern recognition, pages 770–778, 2016.

[34] Hejase, H. A., Mo, Z., Campagna, L., and Siepel, A. 2022. A deep-learning approach for inference of selective sweeps from the ancestral recombination graph. Molecular Biology and Evolution, 39, (1) msab332.

[35] Hermisson, J. and Pennings, P. S. 2005. Soft sweeps: molecular population genetics of adaptation from standing genetic variation. Genetics, 169, (4) 2335–2352.

[36] Isildak, U., Stella, A., and Fumagalli, M. 2021. Distinguishing between recent balancing selection and incomplete sweep using deep neural networks. Molecular Ecology Resources, 21, (8) 2706–2718.

[37] Kelleher, J., Etheridge, A. M., and McVean, G. 2016. Efficient coalescent simulation and genealogical analysis for large sample sizes. PLoS computational biology, 12, (5) e1004842.

[38] Kelleher, J., Thornton, K. R., Ashander, J., and Ralph, P. L. 2018. Efficient pedigree recording for fast population genetics simulation. PLoS computational biology, 14, (11) e1006581.

[39] Kelly, J. K. 1997. A test of neutrality based on interlocus associations. Genetics, 146, (3) 1197–1206.

[40] Kern, A. D. and Schrider, D. R. 2016. Discoal: flexible coalescent simulations with selection. Bioinformatics, 32, (24) 3839–3841.

[41] Kern, A. D. and Schrider, D. R. 2018. diplos/hic: an updated approach to classifying selective sweeps. G3: Genes, Genomes, Genetics, 8, (6) 1959–1970.

[42] Kim, Y. and Nielsen, R. 2004. Linkage disequilibrium as a signature of selective sweeps. Genetics, 167, (3) 1513–1524.

[43] Kingman, J. F. C. 1982. The coalescent. Stochastic processes and their applications, 13, (3) 235–248.

[44] Kipf, T. 2016. Semi-supervised classification with graph convolutional networks. arXiv preprint arXiv:1609.02907.

[45] Korfmann, K., Gaggiotti, O. E., and Fumagalli, M. 2023. Deep learning in population genetics. Genome Biology and Evolution, 15, (2) evad008.

[46] Lauterbur, M. E., Cavassim, M. I. A., Gladstein, A. L., Gower, G., Pope, N. S., Tsambos, G., Adrion, J., Belsare, S., Biddanda, A., Caudill, V., et al. 2023. Expanding the stdpopsim species catalog, and lessons learned for realistic genome simulations. Elife, 12, RP84874.

[47] Lewanski, A. L., Grundler, M. C., and Bradburd, G. S. 2024. The era of the arg: An introduction to ancestral recombination graphs and their significance in empirical evolutionary genomics. Plos Genetics, 20, (1) e1011110.

[48] Li, H. 2011. A new test for detecting recent positive selection that is free from the confounding impacts of demography. Molecular biology and evolution, 28, (1) 365–375.

[49] Lundberg, S. M. and Lee, S.-I. 2017. A unified approach to interpreting model predictions. Advances in neural information processing systems, 30,.

[50] Mo, Z. and Siepel, A. 2023. Domain-adaptive neural networks improve supervised machine learning based on simulated population genetic data. PLoS Genetics, 19, (11) e1011032.

[51] Molnar, C. Interpretable Machine Learning. 2 edition, 2022. URL https://christophm.github.io/interpretable-ml-book/.

[52] Montinaro, F., Pankratov, V., Yelmen, B., Pagani, L., and Mondal, M. 2021. Revisiting the out of africa event with a deep-learning approach. The American Journal of Human Genetics, 108, (11) 2037–2051.

[53] Nait Saada, J., Tsangalidou, Z., Stricker, M., and Palamara, P. F. 2023. Inference of coalescence times and variant ages using convolutional neural networks. Molecular Biology and Evolution, 40, (10) msad211.

[54] Nielsen, R., Vaughn, A. H., and Deng, Y. 2025. Inference and applications of ancestral recombination graphs. Nature Reviews Genetics, 26, (1) 47–58.

[55] Peng, B. and Kimmel, M. 2005. simuPOP: a forward-time population genetics simulation environment. Bioinformatics, 21, (18) 3686–3687.

[56] Perron, L. and Furnon, V. 2019. Or-tools. Google. [Online]. Available: https://developers.google.com/optimization.

[57] Qin, X., Chiang, C. W., and Gaggiotti, O. E. 2022. Deciphering signatures of natural selection via deep learning. Briefings in Bioinformatics, 23, (5) bbac354.

[58] Quelin, A., Austerlitz, F., and Jay, F. 2025. Assessing simulation-based supervised machine learning for demographic parameter inference from genomic data. Heredity, pages 1–10.

[59] Raccuglia, P., Elbert, K. C., Adler, P. D., Falk, C., Wenny, M. B., Mollo, A., Zeller, M., Friedler, S. A., Schrier, J., and Norquist, A. J. 2016. Machine-learning-assisted materials discovery using failed experiments. Nature, 533, (7601) 73–76.

[60] Ray, D. D., Flagel, L., and Schrider, D. R. 2024. Introunet: identifying introgressed alleles via semantic segmentation. PLoS Genetics, 20, (2) e1010657.

[61] Riley, R., Mathieson, I., and Mathieson, S. 2024. Interpreting generative adversarial networks to infer natural selection from genetic data. Genetics, 226, (4) iyae024.

[62] Ronen, R., Udpa, N., Halperin, E., and Bafna, V. 2013. Learning natural selection from the site frequency spectrum. Genetics, 195, (1) 181–193.

[63] Rozemberczki, B., Watson, L., Bayer, P., Yang, H.-T., Kiss, O., Nilsson, S., and Sarkar, R. The shapley value in machine learning. In The 31st International Joint Conference on Artificial Intelligence and the 25th European Conference on Artificial Intelligence, pages 5572–5579. International Joint Conferences on Artificial Intelligence Organization, 2022.

[64] Sanchez, T., Cury, J., Charpiat, G., and Jay, F. July 2020. Deep learning for population size history inference: design, comparison and combination with approximate bayesian computation. Molecular Ecology Resources.

[65] Schrider, D. R. and Kern, A. D. 2015. Inferring selective constraint from population genomic data suggests recent regulatory turnover in the human brain. Genome biology and evolution, 7, (12) 3511–3528.

[66] Schrider, D. R. and Kern, A. D. 2016. S/hic: robust identification of soft and hard sweeps using machine learning. PLoS genetics, 12, (3) e1005928.

[67] Schrider, D. R. and Kern, A. D. 2017. Soft sweeps are the dominant mode of adaptation in the human genome. Molecular biology and evolution, 34, (8) 1863–1877.

[68] Schrider, D. R. and Kern, A. D. 2018. Supervised machine learning for population genetics: a new paradigm. Trends in Genetics, 34, (4) 301–312.

[69] Shapley, L. S. et al. 1953. A value for n-person games.

[70] Sheehan, S. and Song, Y. March 2016. Deep learning for population genetic inference. PLos Computational Biology, 12, (3) 1–28.

[71] Simonyan, K., Vedaldi, A., and Zisserman, A. 2013. Deep inside convolutional networks: Visualising image classification models and saliency maps. arXiv preprint arXiv:1312.6034.

[72] Singh, C., Inala, J. P., Galley, M., Caruana, R., and Gao, J. 2024. Rethinking interpretability in the era of large language models. arXiv preprint arXiv:2402.01761.

[73] Smith, C. C., Tittes, S., Ralph, P. L., and Kern, A. D. 2023. Dispersal inference from population genetic variation using a convolutional neural network. Genetics, 224, (2) iyad068.

[74] Smith, M. L., Ruffley, M., Espíndola, A., Tank, D. C., Sullivan, J., and Carstens, B. C. 2017. Demographic model selection using random forests and the site frequency spectrum. Molecular Ecology, 26, (17) 4562–4573.

[75] Speidel, L., Forest, M., Shi, S., and Myers, S. R. 2019. A method for genome-wide genealogy estimation for thousands of samples. Nature genetics, 51, (9) 1321–1329.

[76] Szatkownik, A., Furtlehner, C., Charpiat, G., Yelmen, B., and Jay, F. 2024. Latent generative modeling of long genetic sequences with gans. bioRxiv, pages 2024–08.

[77] Szatkownik, A., Furtlehner, C., Charpiat, G., Yelmen, B., and Jay, F. Towards creating longer genetic sequences with gans: Generation in principal component space. In Machine Learning in Computational Biology, pages 110–122. PMLR, 2024.

[78] Szatkownik, A., Planche, L., Demeulle, M., Chambe, T., Ávila-Arcos, M. C., Huerta-Sanchez, E., Furtlehner, C., Charpiat, G., Jay, F., and Yelmen, B. 2024. Diffusion-based artificial genomes and their usefulness for local ancestry inference. bioRxiv, pages 2024–10.

[79] Tajima, F. November 1989. Statistical methods for testing the neutral mutation hypothesis by dna polymorphism. Genetics Society of America, 123, (3) 585–595.

[80] Torada, L., Lorenzon, L., Beddis, A., Isildak, U., Pattini, L., Mathieson, S., and Fumagalli, M. 2019. Imagene: a convolutional neural network to quantify natural selection from genomic data. BMC bioinformatics, 20, (Suppl 9) 337.

[81] Tran, L. N., Castellano, D., and Gutenkunst, R. N. 2025. Interpreting supervised machine learning inferences in population genomics using haplotype matrix permutations. Molecular Biology and Evolution, 42, (10) msaf250.

[82] Trost, J., Haag, J., Höhler, D., Jacob, L., Stamatakis, A., and Boussau, B. 2024. Simulations of sequence evolution: how (un) realistic they are and why. Molecular biology and evolution, 41, (1) msad277.

[83] van den Belt, S. and Alachiotis, N. 2025. Fast and accurate deep learning scans for signatures of natural selection in genomes using faster-nn. Communications Biology, 8, (1) 58.

[84] Voight, B. F., Kudaravalli, S., Wen, X., and Pritchard, J. K. 2006. A map of recent positive selection in the human genome. PLoS biology, 4, (3) e72.

[85] Wang, T. and Lin, Q. 2021. Hybrid predictive models: When an interpretable model collaborates with a black-box model. Journal of Machine Learning Research, 22, (137) 1–38.

[86] Wang, Z., Wang, J., Kourakos, M., Hoang, N., Lee, H. H., Mathieson, I., and Mathieson, S. 2021. Automatic inference of demographic parameters using generative adversarial networks. Molecular ecology resources, 21, (8) 2689–2705.

[87] Whitehouse, L. S. and Schrider, D. R. 2023. Timesweeper: accurately identifying selective sweeps using population genomic time series. Genetics, 224, (3) iyad084.

[88] Whitehouse, L. S., Ray, D. D., and Schrider, D. R. 2024. Tree sequences as a general-purpose tool for population genetic inference. Molecular Biology and Evolution, 41, (11) msae223.

[89] Wright, F. A., Huang, H., Guan, X., Gamiel, K., Jeffries, C., Barry, W. T., Pardo-Manuel de Villena, F., Sullivan, P. F., Wilhelmsen, K. C., and Zou, F. 2007. Simulating association studies: a data-based resampling method for candidate regions or whole genome scans. Bioinformatics, 23, (19) 2581–2588.

[90] Yelmen, B. and Jay, F. 2023. An overview of deep generative models in functional and evolutionary genomics. Annual Review of Biomedical Data Science, 6, (1) 173–189.

[91] Yelmen, B., Decelle, A., Ongaro, L., Marnetto, D., Tallec, C., Montinaro, F., Furtlehner, C., Pagani, L., and Jay, F. 2021. Creating artificial human genomes using generative neural networks. PLoS genetics, 17, (2) e1009303.

[92] Yelmen, B., Decelle, A., Boulos, L. L., Szatkownik, A., Furtlehner, C., Charpiat, G., and Jay, F. 2023. Deep convolutional and conditional neural networks for large-scale genomic data generation. PLOS Computational Biology, 19, (10) e1011584.

[93] Zhang, Q.-s. and Zhu, S.-C. 2018. Visual interpretability for deep learning: a survey. Frontiers of Information Technology & Electronic Engineering, 19, (1) 27–39.

[94] Zhang, Q., Yang, Y., Ma, H., and Wu, Y. N. Interpreting cnns via decision trees. In Proceedings of the IEEE/CVF conference on computer vision and pattern recognition, pages 6261–6270, 2019.

