## Supplementary Material for "Interpreting Convolutional Neural Networks in Population Genetics"

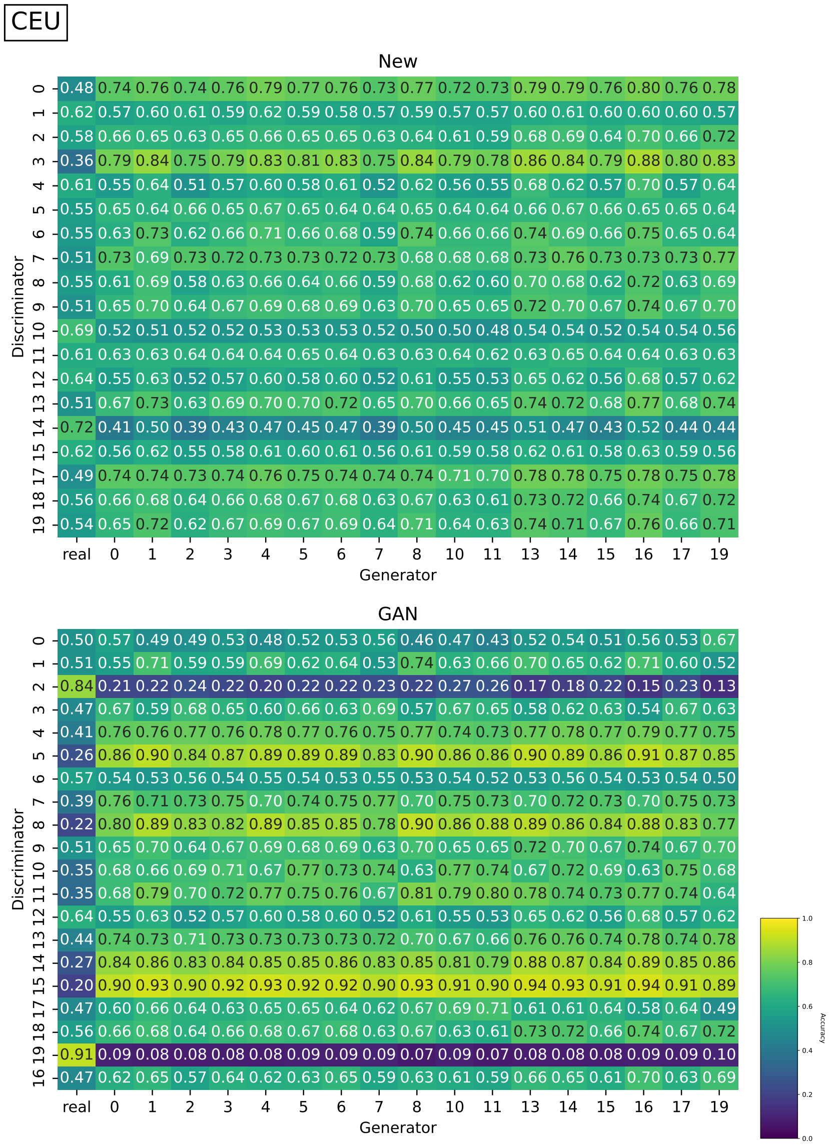


Figure S1: Accuracies of newly trained CNNs (top) and `pg-gan` discriminator CNNs (bottom) for the CEU population. Results are separated into real (column 1 of each heatmap) and generated data, and further separated into simulated data from distinct pg-gan generators. Generators are filtered to only those which did not collapse during GAN training. Discriminators are filtered out which predicted the same value for every input.

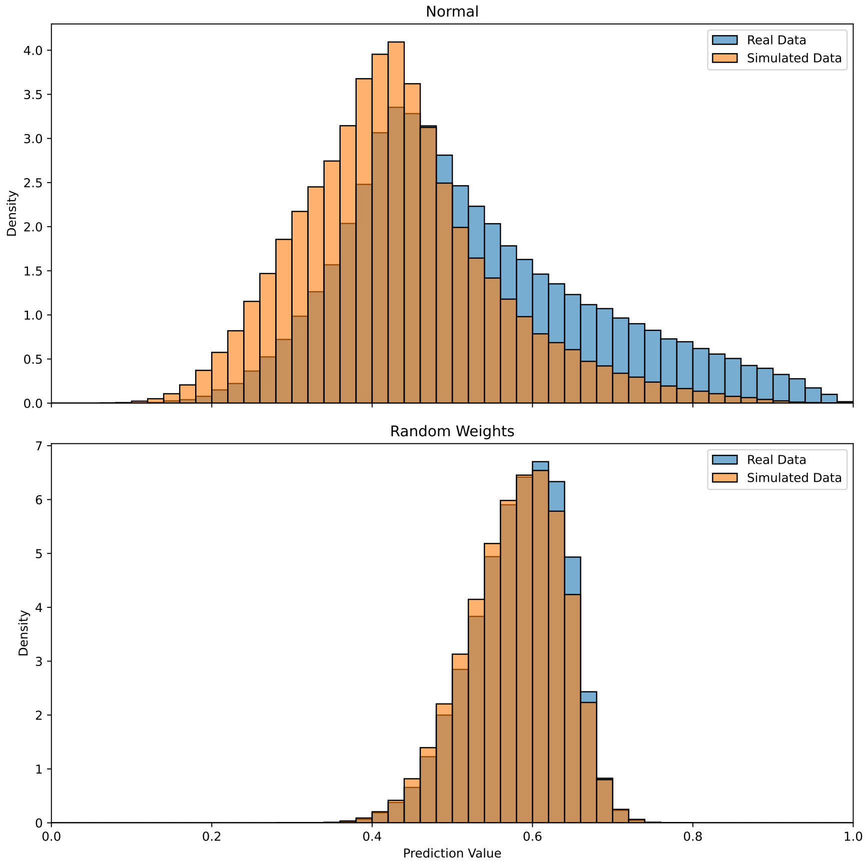


Figure S2: Predictions (probabilities of real) for retrained CNN discriminator seed 0 (“Normal”) vs. a randomly initialized CNN (“Random Weights”). For the random weights, the prediction values are closer to 0.5, with less separation between real and simulated data. For the retrained network the predictions are more spread out, with more separation between real and simulated data. However this is not a bimodal distribution, indicating the task of distinguishing between the data source is difficult.

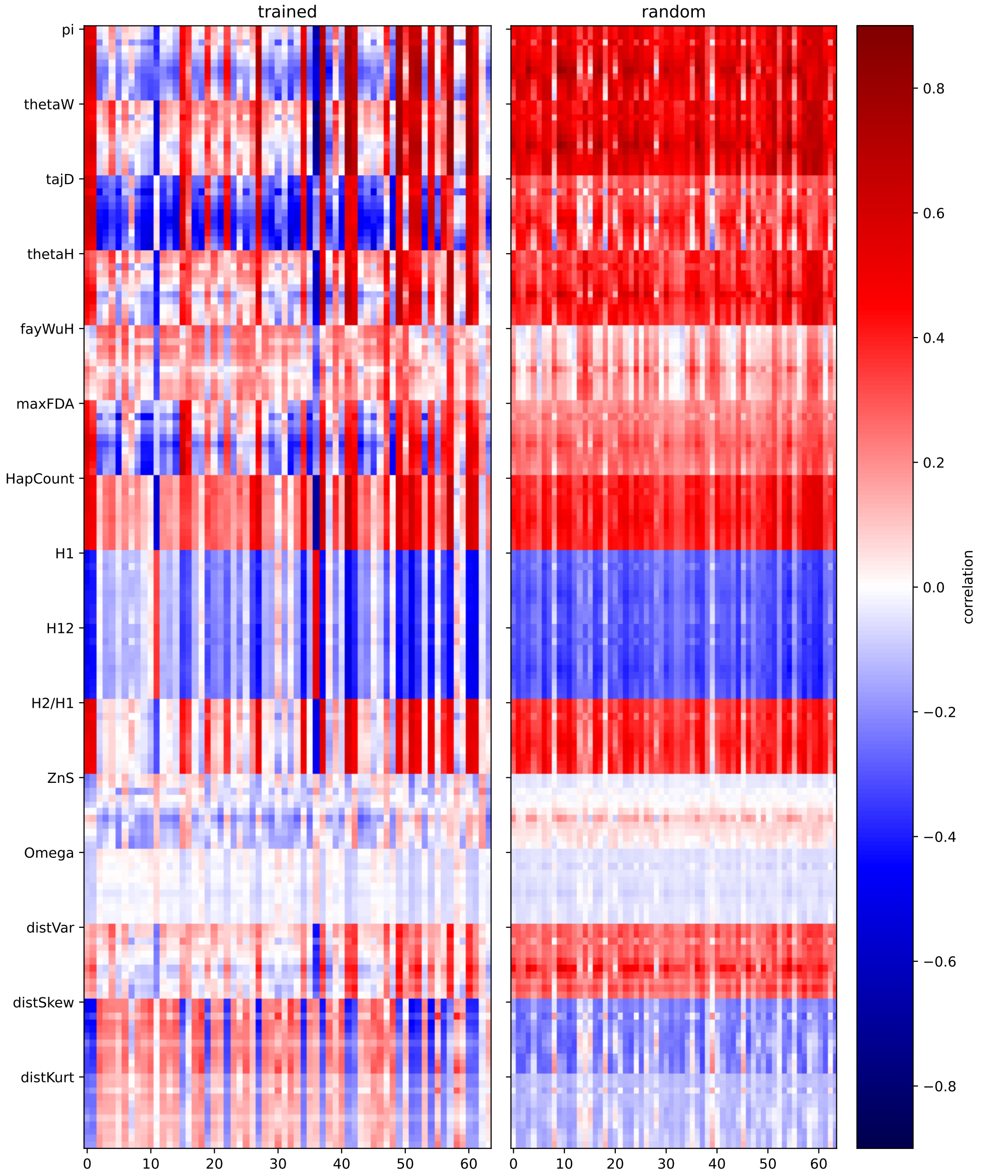


Figure S3: Correlation between last hidden layer values and summary statistics for a trained ResNet CNN for sweep detection (left) vs. a randomly initialized network with the same architecture (right). The $x$-axis shows the 64 features of the last hidden layer, clustered by similarity of their correlation profiles. The $y$-axis shows each of `diploSHIC`'s summary statistics, which are measured in 11 adjacent windows spanning the simulated region.

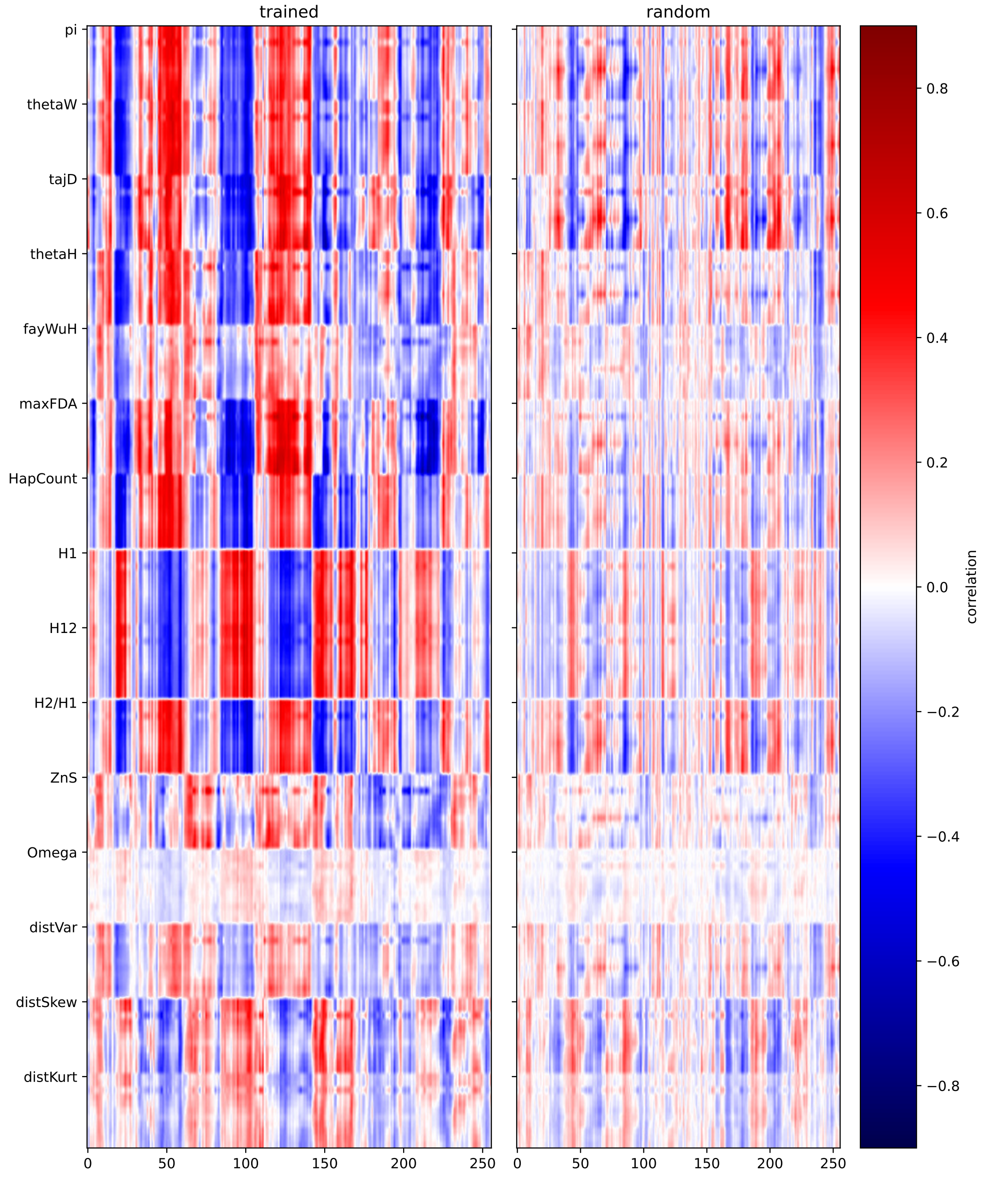


Figure S4: Side by side comparison of correlations of raw GCN features with `diploSHIC`'s summary stats. For a pre-trained network on the left and random weights on the right.

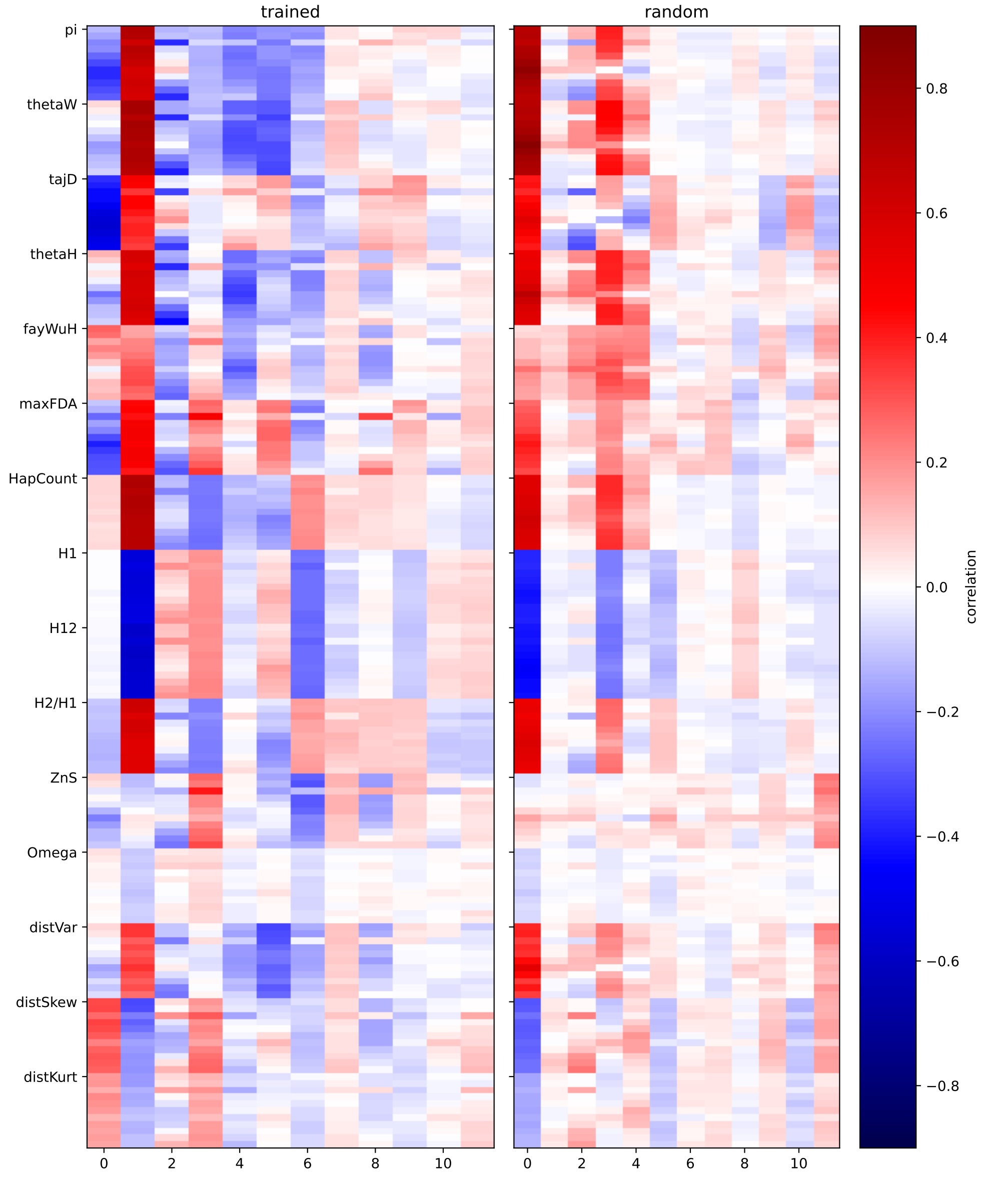


Figure S5: Side by side comparison of correlations of transformed ResNet features (first 12 principal components) with diploShic summary stats. For a pre-trained network on the left and randomly initialized weights on the right.

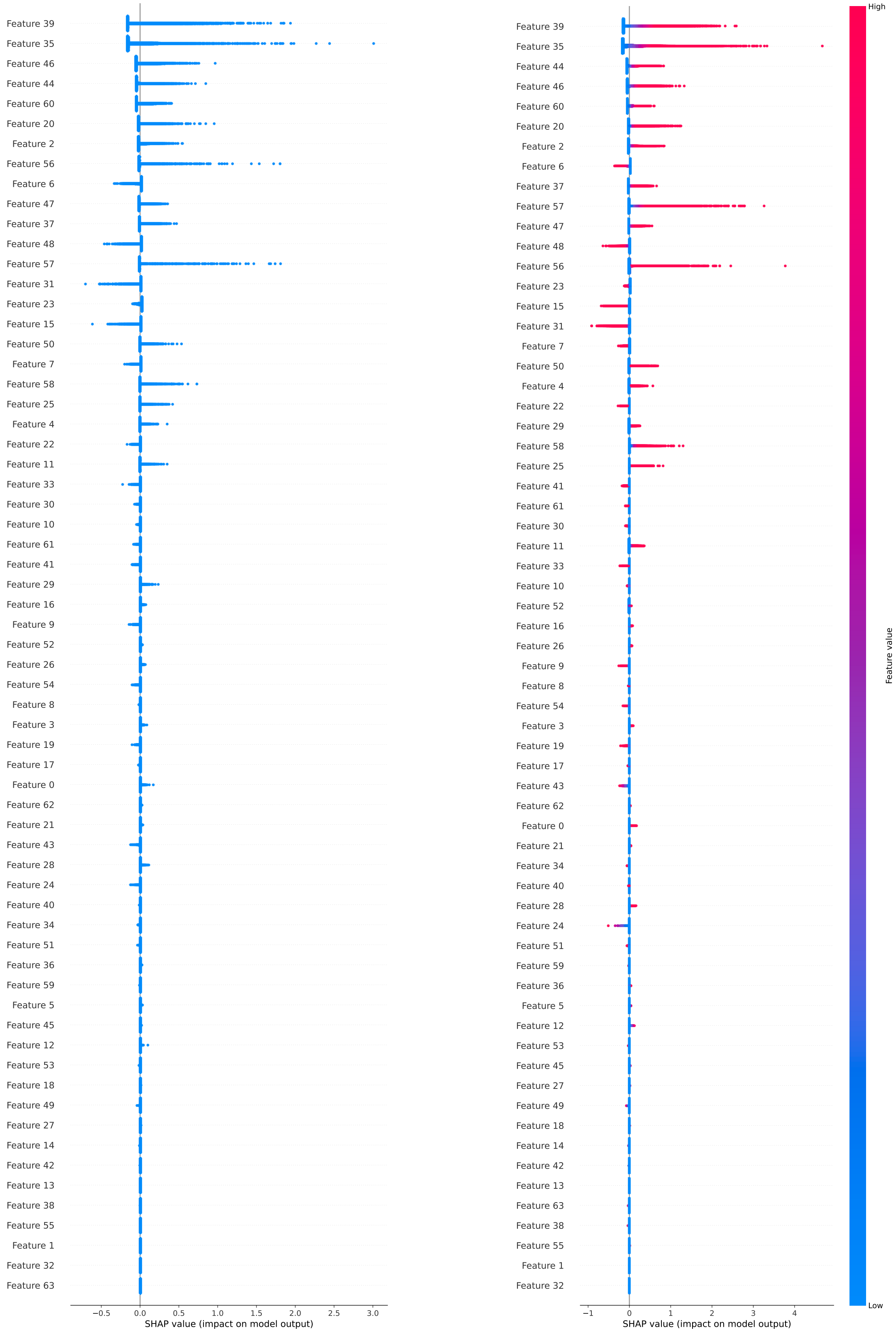


Figure S6: SHAP values for learned features for seed 0 (CEU). Left: Deep SHAP values. Right: Permutation SHAP values.
